# A multiscale whole-cell theory for mechano-sensitive migration on viscoelastic substrates

**DOI:** 10.1101/2022.04.01.486783

**Authors:** Wenya Shu, C. Nadir Kaplan

## Abstract

Increasing experimental evidence validates that both the elastic stiffness and viscosity of the extracellular matrix regulate mesenchymal cell behavior, such as the rational switch between durotaxis (cell migration to stiffer regions), anti-durotaxis (migration to softer regions), and adurotaxis (stiffness-insensitive migration). To reveal the mechanisms underlying the crossover between these motility regimes, we have developed a multiscale chemo-mechanical whole-cell theory for mesenchymal migration. Our framework couples the sub-cellular focal adhesion dynamics at the cell-substrate interface with the cellular cytoskeletal mechanics and the chemical signaling pathways involving Rho GTPase proteins. Upon polarization by the Rho GTPase gradients, our simulated cell migrates by concerted peripheral protrusions and contractions, a hallmark of the mesenchymal mode. The resulting cell dynamics quantitatively reproduces the experimental migration speed as a function of the uniform substrate stiffness and explains the influence of viscosity on the migration efficiency. In the presence of stiffness gradients and absence of chemical polarization, our simulated cell can exhibit durotaxis, anti-durotaxis, and adurotaxis respectively with increasing substrate stiffness or viscosity. The cell moves toward an optimally stiff region from softer regions during durotaxis and from stiffer regions during anti-durotaxis. We show that cell polarization through steep Rho GTPase gradients can reverse the migration direction dictated by the mechanical cues. Overall, our theory demonstrates that opposing durotactic behaviors emerge via the interplay between intracellular signaling and cell-medium mechanical interactions in agreement with experiments, thereby elucidating complex mechano-sensing at the single-cell level.

## Introduction

Mesenchymal migration is fundamental to physiological and pathological phenomena such as development, growth, and tumor invasion. A mesenchymal cell moves by means of the recurring formation and removal of peripheral focal adhesion (FA) sites where it senses the mechanical properties of the extracellular matrix (ECM) [1–5]. At each FA site, the cell spreads at a speed set by the competition between actin polymerization, actomyosin contractility due to the pulling of the myosin motors, and mechanical stresses accross the cell-ECM interface. The net migration speed is determined at the whole-cell level by the contributions from all FA sites, highlighting the multiscale nature of the process, and has a nonmonotonic dependence on the ECM elastic stiffness [6–9]. This trend was qualitatively explained by a stochastic theory of the “motor-clutch” force transmission on a single FA site at the cell anterior [10, 11]. Because biological tissues are viscoelastic, the motor-clutch model was recently extended to investigate the cellular response to viscoelasticity [12–14].

Despite the successes of the single FA theories, the physico-chemical dynamics of motility requires a multi-scale whole-cell description. This is because, first, single FA models inherently overestimate the measured migration speeds since the opposing traction forces across the distant FA sites of a cell are ignored. Second, cells must detect the substrate heterogeneity with a higher resolution across their body than at the single FA scale. For example, vascular smooth muscle cells, glioblastoma cells (U-87MG and U-251MG), and MDA-MB-231 breast cancer cells can move up the stiffness gradients, termed durotaxis [15–18]. MDA-MB-231 and U-251MG can also migrate against stiffness gradients, known as anti-durotaxis, or become adurotactic by being insensitive to the gradients [17–19]. Some theories replicated durotaxis by taking the traction force or migration speed as explicit functions of substrate stiffness [20, 21]. However, since durotaxis is *a posteriori* built into these models, they cannot address the switch between durotaxis and anti-durotaxis. Third, migration is regulated by the cell-level reaction-transport dynamics of an intracellular biopolymer network. This involves the two prominent GTPase proteins, such as Rac1, which controls actin polymerization and focal complex assembly, and RhoA, which modulates actomyosin contractility [22–24]. A multi-scale approach that considers the biochemical signaling dynamics together with the responses of distinct FA sites to the local substrate mechanics could potentially explain the experimental migration speeds and the gradient sensitivity.

Here, we present a multiscale chemo-mechanical whole-cell theory for mesenchymal migration on viscoelastic substrates. Our model couples the sub-cellular mechanics of the substrate–FA site interfaces with the cell-scale mechanics of the nucleus–cytoskeletal network complex and reaction-diffusion dynamics of the Rac1 and RhoA GTPase proteins. Hence, our framework differentiates itself from the existing whole-cell theories that are purely mechanical or do not explicitly incorporate the sub-cellular FA dynamics [25–30]. Our simulated cell can acquire a chemically induced polarization and a dynamic complex morphology during migration, with a speed profile that is biphasic with respect to the substrate elasticity about a threshold stiffness, in quantitative agreement with experiments. Moreover, for a Kelvin-Voigt viscoelastic substrate, we analytically determine the traction force at an FA site in terms of the substrate viscosity and stiffness, and explain how the migration speed changes biphasically about a threshold viscosity, beyond which the cell abruptly loses traction. Importantly, the explicit treatment of the cell FA sites allows us to investigate the migration on nonuniform substrates with stiffness gradients. For a chemically apolar cell, we identify the crossover between durotaxis, anti-durotaxis, and adurotaxis as a function of the substrate stiffness and viscosity, and propose a unifying mechanism for the mechanosensitivity of mesenchymal migration. We further show that switching on the chemical polarity can override the cell mechanical response and reverse the migration direction. Altogether, our results explain how the observed cell motility patterns are governed by the interplay between cell signaling, substrate elasticity and viscosity, which are difficult to independently control in experiments.

## Multiscale chemo-mechanical theory for mesenchymal migration

The biological components included in our multiscale whole-cell theory are illustrated in Fig. 1 A, B. Mesenchymal migration is initiated by an emergent intracellular polarization that drives the formation of leading edges with actin-rich protrusion sites and the actomyosin-induced retraction at the cell rear. Upon polarization, FA sites form at the protruding edges. Each vertex in our model is assigned a FA site (Fig. 1 C, D). The polarization is induced (i) by the gradients in the substrate material properties, which require modeling the mechanical cell-substrate interactions, and/or (ii) by the distribution of Rho GTPase proteins in inactive or active states across the cell, which necessitate a continuum description in the form of cell-level reaction-diffusion equations [22, 27, 29]. At the sub-cellular level, coupling (i) and (ii) with a motor-clutch model for the FA sites at each vertex allows us to determine how the Rho GTPases regulate actin polymerization, tensile actomyosin forces, and adhesion dynamics (Fig. 1 B, C). At the cell level, the forces generated at the FA sites, the membrane resistance forces, the cytoskeletal elastic forces, and the cytoplasmic viscous forces are all transferred to the nucleus, which serves as the hub for global force transmission across the cell in our model (Fig. 1 D, E). For the Rho GTPase dynamics, we adopt the reaction-diffusion framework introduced in Ref. [29]. To model each FA site, we generalize the motor-clutch model for a viscoelastic substrate in Ref. [14] to include the effect of the Rho GTPases and the substrate heterogeneity. We neglect the inertial forces since cells migrate in low-Reynolds-number environments. In our formulation, all variables are time dependent, the time derivative of a variable *f* = *f*(*t*) is denoted by 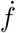, and *f* ^*i*^ represents the value of *f* at the vertex *i* across the model cell.

**Fig. 1:**
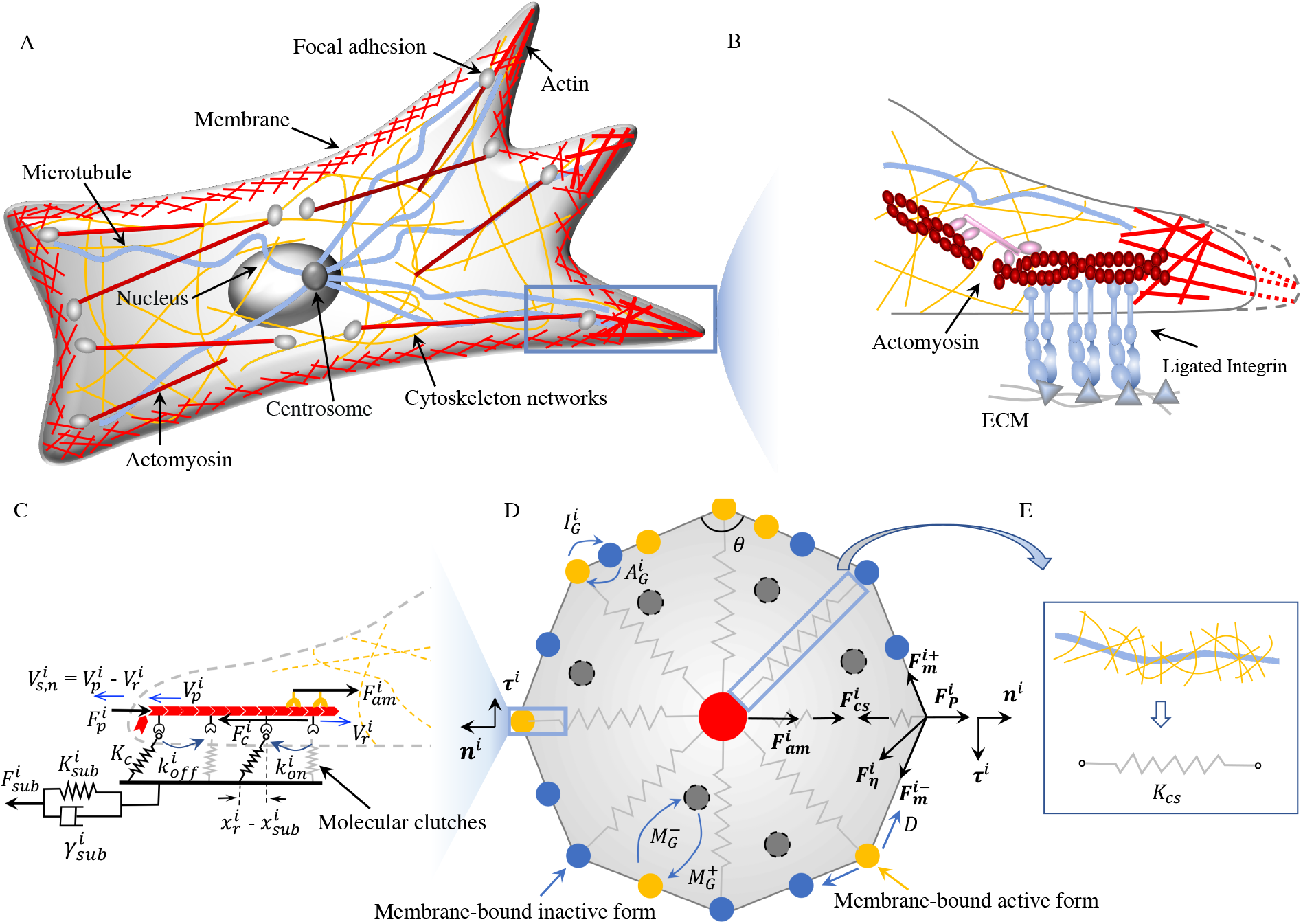
Whole-cell model schematics. (A) Intracellular organization that drives nucleus movement and mesenchymal migration. (B) Close-up view of the cell edge depicts the coupling between the actomyosin complexes with myosin motors, molecular clutches (blue) between the cell and the integrin ligands of the extracellular matrix (ECM) at the focal adhesion site, and polymerizing actin (red lines) that drive edge protrusion. The spreading velocity at vertex *i* is 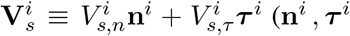: unit radial and polar vectors). (C) Positions, speeds, forces, and material parameters underlying the sub-cellular level equations. (D) The cell-level vertex model. The chemical signaling is mediated by the Rac1 (*R*) and RhoA (*ρ*) GTPase proteins in the membrane-bound active (yellow dots) and inactive states (blue dots), and inactive state dissolved in the cytoplasm (gray dots). The conversion rates between the three states are 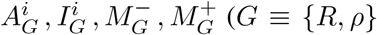; see Eq. 12). (E) The elastic forces on cytoskeletal biopolymers (microtubules and intermediate filaments) are modeled by Hookean springs with the stiffness *K*_*cs*_.

### Equations of motion at sub-cellular level

Mesenchymal cell migration is maintained by the peripheral FA sites where molecular clutches are formed temporarily between the integrin ligands bound to the viscoelastic ECM and the intracellular actomyosin complex (Fig. 1 B, C). At the sub-cellular level of our model, each vertex *i* comprises an FA site that transmits forces between the ECM and actomyosin, and an elastic membrane section (Fig. 1 C, D). A cytoskeletal “spring” linked to a vertex represents the microtubules and intermediate filaments, which transmit forces between the nucleus, the FA site, and the membrane (Fig. 1 D, E). Denoting **x**^*i*^(*t*) as the position vector of the vertex *i*, we characterize the edge protrusions and retractions with a local spreading velocity 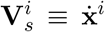, which is regulated by the Rho GTPase proteins. At vertex *i*, we consider the two prominent Rho GTPase members Rac1 (*R*^*i*^) and RhoA (*ρ*^*i*^); each can be in 2 membrane-bound states with active concentrations 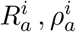 and inactive concentrations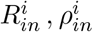. For simplicity, our model assumes that active chemo-mechanical processes determine solely the radial spreading speed along the cytoskeletal springs 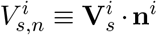 whereas the polar spreading speed 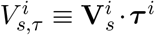 is set by the passive membrane deformations (**n**^*i*^, ***τ*** ^*i*^ defined in Fig. 1 D).

The radial spreading speed 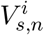 is dictated by the competition between actin polimerization, myosin motor forces, and the membrane resistance. Actin filaments polymerize against the cell membrane with a polymerization speed 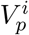at each vertex *i* (Fig.1 B, C). In our model, 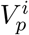 is determined by the active Rac1 concentration 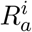 and its mean 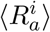 averaged over all vertices as 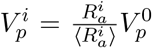, where 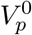 is the reference polymerization speed. The inward pulling forces of the myosin motors and the membrane elastic resistance counteract actin polymerization by inducing an actin retrograde flow with a speed 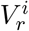 from the edge to the interior of the cell. Then, 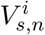 is given by

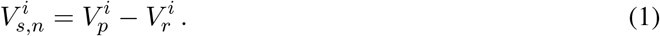

The retrograde flow speed 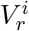 is governed by the force balance at the FA sites and will be detailed below. Eq. 1 links the spreading dynamics to the local active Rac1 GTPase concentration and its cell-level average, building in a coupling across the two scales.

The actin retrograde flow is actively controlled by the myosin motors within the peripheral actomyosin complex. At the FA site where the cell forms molecular clutches between actomyosin and the ECM, the retrograde flow speed is defined as 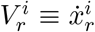 where 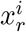 is displacement of the actomyosin end of the FA site at vertex *i* (Fig. 1 C). Defining 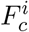 as the elastic restoring force of the bounded clutches and 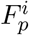 as the net force resisting protrusion, the net local pulling force acting on the actomyosin complex

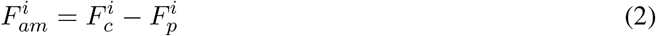

regulates 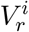 experimentally according to the empirical force-velocity relation [10]

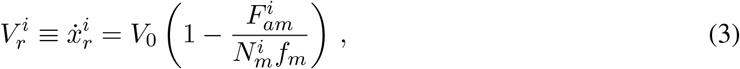

where *V*_0_ is the unloaded myosin motor speed at 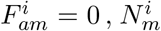 is the number of active myosin motors, and *f*_*m*_ is the force that stalls the activity of one myosin motor. Because active RhoA proteins enhance myosin-induced cell contraction [31, 32], we relate 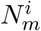 to the active RhoA concentration 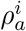 as 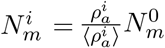, where 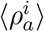 is the mean active RhoA concentration averaged over all vertices and 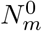 is the reference myosin motor number. In Eq. 2, 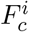 is set by the cell-medium interactions at equilibrium at the FA site, and 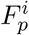 is computed from a force balance across the vertex *i* and the attached cytoskeletal elements (Fig. 1 D, E). We determine these two forces next.

At each FA site, the bound clutches between the extracellular integrin ligands and the intracellular actomyosin complex must generate a restoring force between the displacement at the filament end 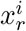 and that at the substrate end 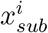 (Fig. 1 C). Defining *K*_*c*_ as the clutch spring constant, the restoring force on a single clutch is 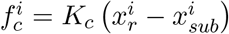. Denoting the number of clutches as 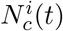 and ≤ *P* ^*i*^(*t*) ≤ 1 as the fraction of the bound clutches, the total restoring clutch force at vertex *i* is given by

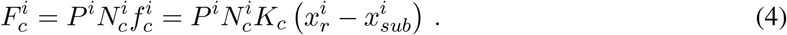

Since Rac1 proteins are required for the focal complex assembly [33–35], we assume that 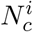 is dynamically controlled by the local Rac1 concentration, i.e., 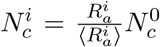, where 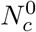 denotes the reference clutch number. With an association rate 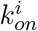 and the dissociation rate 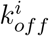, the time evolution of *P* ^*i*^ in Eq. 4 is governed by the mean-field rate equation [13, 14, 36]

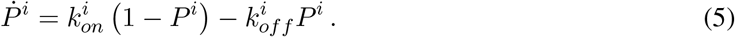

In Eq. 5, we assume that the bound clutches dissociate with a catch-bond behavior with the rate 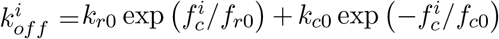. Here, *k*_*r*0_ and *k*_*c*0_ denote the unloaded off-rate and the unloaded catch-rate, respectively, *f*_*r*0_ is the characteristic rupture force, and *f*_*c*0_ the characteristic catch force.

At mechanical equilibrium across the FA site, the elastic force given by Eq. 4 must be balanced by the viscoelastic deformation force of the substrate with a strength 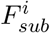. Defining the local elastic stiffness as 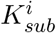 and uniform substrate viscosity as *γ*_*sub*_, we use the Kelvin-Voigt model 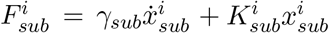. (Fig. 1 C) [12]. Then, using Eq. 4 and the equality 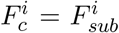 yield the equation of motion for the displacement 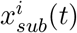 at the subtrate end of the focal adhesion as

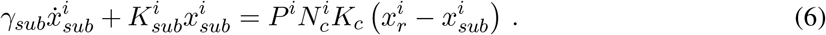

The dependence of 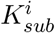 on the vertex label *i* will allow us to examine the effect of the substrate stiffness gradients on the migration dynamics.

The net peripheral force resisting protrusion 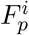 in Eq. 2 is determined by the balance between the radial forces on the cytoskeletal elements and those at the cell boundary (Fig.1 C, D). The cytoskeletal elements consisting of microtubules and intermediate filaments link the vertices to the cell nucleus and are represented by a Hookean spring with a stiffness constant *K*_*cs*_ in our model (Fig.1E). Denoting the position vector of the cell nucleus by **x**_*nuc*_(*t*), the distance between the cell nucleus and the *i*^*th*^ vertex of the cell is is given by the radius *r*_0_ in the undeformed (initial circular) configuration and by *r*^*i*^ ≡ |(**x**_*nuc*_ − **x**^*i*^)| in the deformed configuration. The radial restoring force is then expressed as

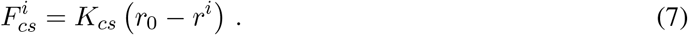

Representing the cell membrane by a Hookean spring with stiffness *K*_*m*_ between two adjacent vertices, the tensile membrane forces at the *i*^*th*^ vertex are given by 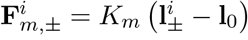, where 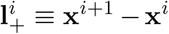 and 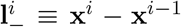 denote the distance vectors between the vertex *i* and its two neighbors, and **l**_0_ is the equilibrium initial distance vector between two adjacent vertices. Defining the deformed membrane length about the *i*^*th*^ vertex as 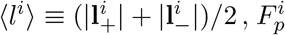 is obtained from the radial force balance

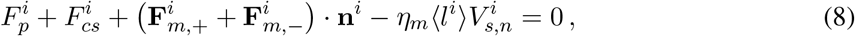

where the last term on the left-hand side denotes the viscous drag with the viscosity *η*_*m*_ due to interactions between the cell membrane and the surrounding medium.

The vertex position can be computed when, in addition to the radial spreading speed, the polar spreading speed 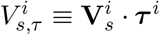 in response to the passive membrane deformations is obtained by

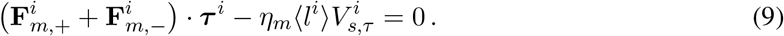

Eqs. 1– 9 fully determine the vertex spreading velocity 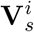, provided that the nucleus position **x**_*nuc*_ and the membrane-bound Rho GTPase concentrations 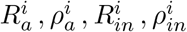 at all vertices are known. We will specify **x**_*nuc*_(*t*) from cell-level force balance and 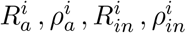 from a set of reaction-diffusion equations coupled with the dynamics of Rho GTPase proteins dissolved in the cytoplasm.

### Equations of motion at cellular level

As the largest and stiffest organelle, the nucleus transmits forces between different sets of cytoskeletal microtubule and intermediate filament bundles, where each set is linked to a separate FA site and a membrane section (Fig. 1 D) [37]. The resulting cell-level network maintains the structural integrity of the cell by imposing a global force balance, which governs the nucleus translocation mostly independent of the centrosome position [38]. That is, the sum of the subcellular forces at each cytokeletal spring attached to the vertex *i*, i.e., the active force on the actomyosin stress fiber 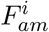 (Eq. 2) and the passive cytoskeletal spring force 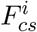 (Eq. 7), determines the net nuclear force **F**_*nuc*_(*t*) according to the equilibrium condition (Fig. 1 D) (*N* : total number of vertices),

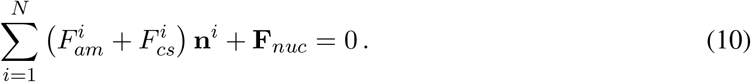

The nucleus position **x**_*nuc*_ is governed by **F**_*nuc*_ noting that the nucleus undergoes passive drag with a velocity 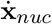 in the viscous cytoplasm. Approximating the nucleus to a rigid sphere with radius *r*_*nuc*_ and denoting the cytoplasm viscosity by *η*_*cp*_, the viscous drag force is expressed by the Stokes’ law as [26]

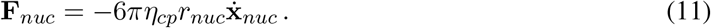

Eqs. 10 and 11 complement Eqs. 1– 9 to determine the motion of the nucleus and all vertices simultaneously, thereby closing the mechanical feedback loop between the sub-cellular-and cell-level dynamics.

The distributions of the membrane-bound active Rac1 and RhoA GTPase concentrations 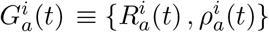 at the cell level are set by three main mechanisms: First, they can diffuse between adjacent vertices; second, they can transition to/from the membrane-bound inactive concentrations 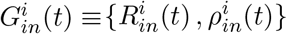, third, there can be a turnover between 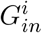 and the cytoplasmic concentrations 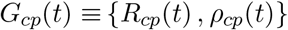. To combine these three mechanisms, we adopt the biochemical reaction-diffusion model introduced in Ref. [29]. We further assume that *G*_*cp*_ remain spatially uniform in the cytoplasm at all times, and the total Rac1 and RhoA concentrations are constant over the course of migration. As depicted in Fig. 1 D, inactive-to-active conversion rates are 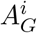, and active-to-inactive conversion rates are 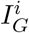 where *G* ≡ {*R, ρ*}. The exchange between the membrane-bound inactive protein concentrations and the cytoplasmic concentrations takes place with an association rate 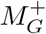 and disassociation rate 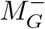.

The diffusive fluxes of active and inactive proteins between adjacent vertices are discretized into finite difference terms. The reaction-diffusion equations are then given by (*y* ≡ {*a, in*}),

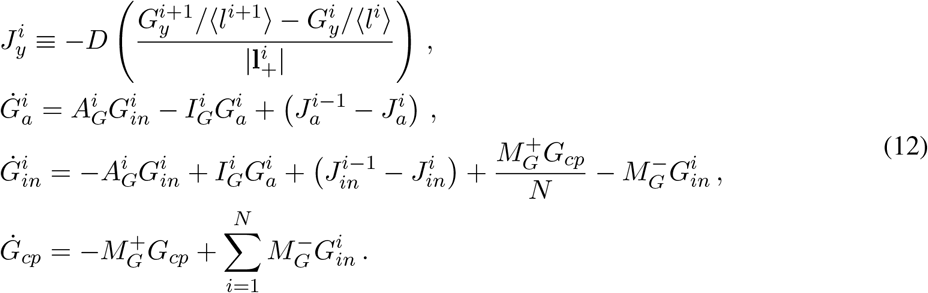

The local active Rac1 and RhoA GTPase concentrations 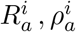 and their averages over all vertices 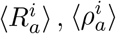 control the cell deformations by regulating the actin polymerization speed 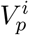 Eq. 1), the number of active myosin motors 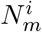 (Eq. 3), and the clutch number 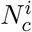 an FA site (Eq. 4). To incorporate reverse coupling from the membrane deformations to the rate terms, the self-reinforcements of 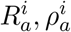 and the competition in-between, we assume the functional forms 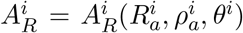 in Eq. S1 and 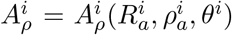 in Eq. S2 (Sec. S1). Here, *θ*^*i*^ is the angle between the two membrane sections at each vertex (Fig. 1 D). Eqs. S1 and S2 couple 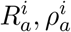 for recurrent polarization and the recovery of cell deformations. Particularly, an excessive increase in *θ*^*i*^ at the retracting cell rear builds up 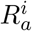 to increase actin polymerization and thus reverse the retraction at the end of each migration step. A large decrease in *θ*^*i*^ at the protruding cell front boosts *ρ*_*a*_ that increases actomyosin contraction to resist further protrusion.

## Results

By numerically solving the two-level whole-cell theory, we investigate the emergent polarity and complex morphology of a mesenchymal cell, as well as its migration dynamics as a function of the substrate elasticity, viscosity, and the gradients thereof. Due to the discrete nature of our vertex model, Eqs. 1– 9 for the vertex spreading velocity 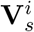, Eqs. 10, 11 for the nucleus position **x**_*nuc*_, and Eqs. 12, S1, S2 for the Rho GTPase concentrations in the local active state 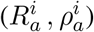, local inactive state 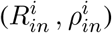, and global cytoplasmic state (*R*_*cp*_, *ρ*_*in*_) constitute a set of nonlinear ordinary differential equations in time *t*. The equations are non-dimensionalized by the position scale *X* ≡2*πr*_0_ ≈30 *µm*, speed scale *V*_0_ = 120 *nm/s*, time scale *T X/V* 260 *s*, and force scale 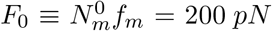 (Table S1). To solve this initial value problem, we approximate an initially circular cell by an *N*-sided polygon with its corners setting the *N* vertex positions **x**^*i*^ at *t* = 0. All initial forces, velocities, displacements, and *P* ^*i*^ (Eq. 4) are set to 0. Uniform or random initial conditions for 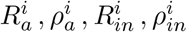 are specified sep-arately for each study and will be given at each section. The total concentrations of Rac1 and RhoA are each conserved, which set *R*_*cp*_, *ρ*_*cp*_ at *t* = 0 (Eq. S4). Starting from these initial conditions, the time evolution of the model cell is simulated according to the algorithm presented in Fig. S1. When the cytoplasmic inactive Rac1 concentration *R*_*cp*_ reaches steady state, we reinitialize 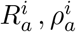 to ensure that migration continues (see Fig. S1). Each simulation considers *Q* > 10 such cycles of the Rho GTPase relaxation dynamics. For the fixed experimental parameters listed in Table S2, we characterize the effect of substrate viscoelasticity on mesenchymal migration for an experimentally relevant range of the ECM stiffness and viscosity (Table 1). On nonuniform substrates with prescribed stiffness gradients, each vertex is assigned a local stiffness 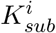 depending on its position on the substrate at each time step, whereas on uniform substrates, 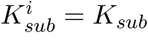 for all *i* = {1, …, *N*} at all times. A mesh convergence study that we have performed demonstrates that *N* = 16 − 24 yield good numerical accuracy (Sec. S2). Therefore, we choose *N* = 16 to lower the computational cost of the simulations. Each simulation involves ∼10^6^ time steps with a time step Δ*t* = 0.002*s* (Δ*t* ≈10^−5^ in dimensionless units) corresponding to a migration dynamics over at least an hour in real units.

**Table 1:**
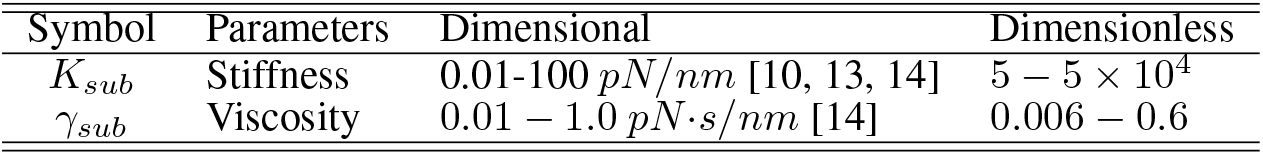
Substrate stiffness and viscosity ranges used in the simulations.

### Chemically induced cell polarity steers migration with complex morphology

Cell polarization followed by directional locomotion on a uniform substrate can be driven by a nonuniform distribution of the Rho GTPase proteins. To simulate this scenario, we set a random initial distribution of the active and inactive Rac1 and RhoA concentrations on all vertices except that two randomly selected vertices of the rightmost five vertices in Fig.2 A have a higher active Rac1 concentration 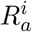 at *t* = 0 (Section S3, Fig. S3). Accordingly, at the vertex *i* = *A*, a high local 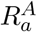 enhances the actin polymerization speed 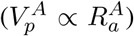 and also the number of molecular clutches 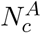, in turn producing a strong focal adhesion evidenced by a high substrate traction force 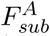 as per Eq. 6 (Fig.2 A, B). Strong adhesion effectively suppresses the actin retrograde flow speed 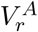 and generates a ‘sticking’ state of the actin filaments. This speeds up actin polymerization (i.e., high 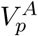 ; see Eq. 1) to protrude the cell membrane at the vertex A (Fig. 2 A, *t/T* = 0.12). In contrast, low 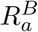 at the vertex *B* gives rise to low 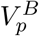 and low 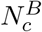.

**Fig. 2:**
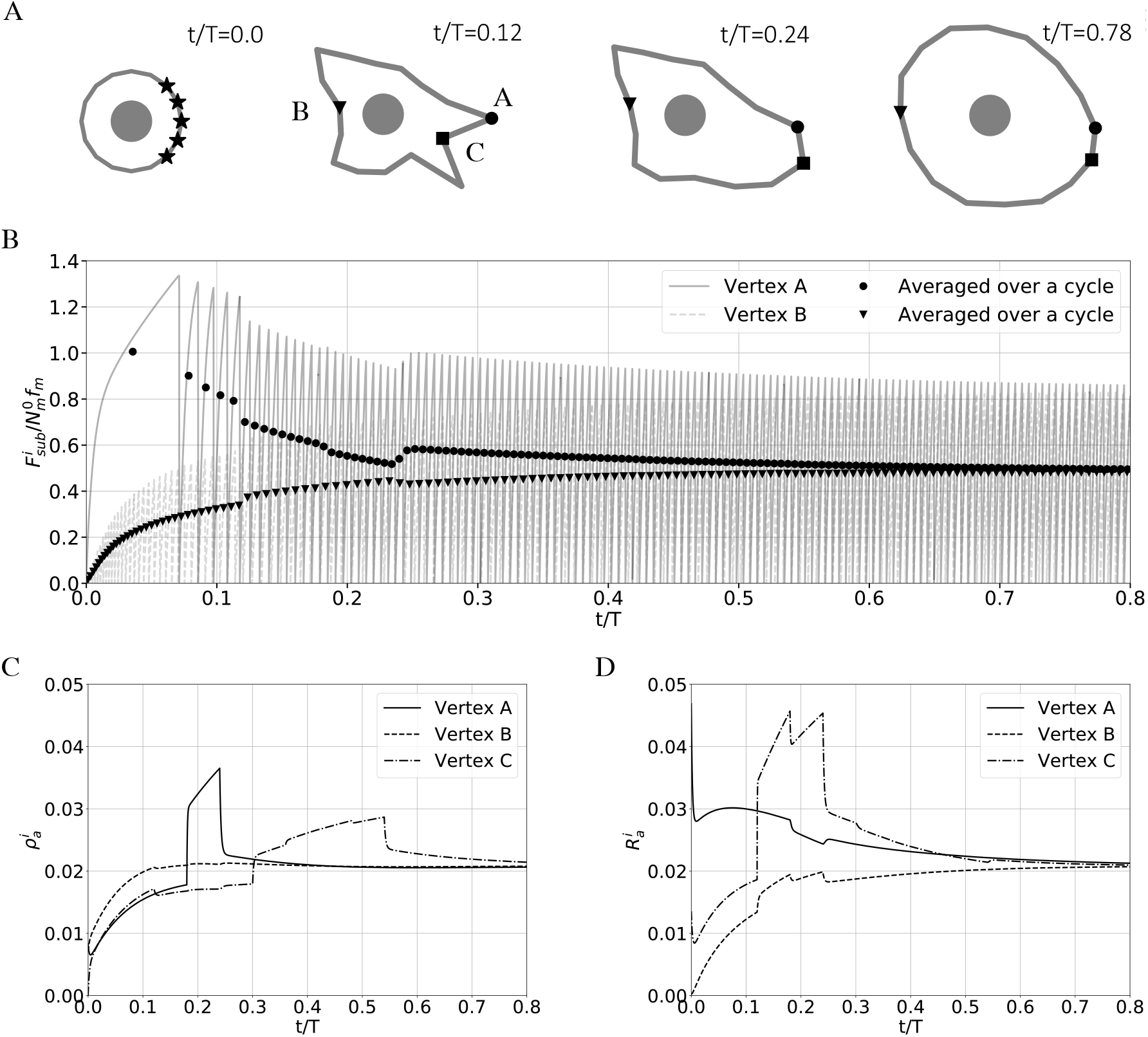
Chemically induced cell polarity and migration. (A) A migrating cell with polarity induced shape change at four different times. Three vertices with strong protrusion or contraction dynamics are labeled as *A, B, C*. At time *t* = 0, our simulation algorithm randomly assigns a strong active Rac1 signal to one or two vertices from the five rightmost vertices denoted by stars (see Fig. S3 A). (B) Over one Rho GTPase relaxation cycle (*Q* = 1), time series of the dimensionless substrate force at the leading vertex *A* (full line) and the trailing vertex *B* (dashed line). The dots and inverse triangles denote the average values over a single period of clutch formation and breakup. Time evolutions of the (C) membrane-bound active RhoA concentration 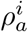 and (D) active Rac1 concentration 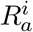 on *A, B, C*.

Correspondingly, the focal adhesion at the vertex *B* allows for a high 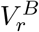, sustaining a much lower 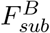 (Fig. 2 B). Such a ‘slipping state’ of the actin filaments coupled with weak actin polymerization retracts the vertex *B* (Fig.2 A, *t* = 0.12). The coexistence of the sticking state with a slow retrograde flow at *A* and the slipping state with a fast retrograde flow at *B* breaks the cell symmetry [39]. Taken together, a polarized distribution of the Rac1 proteins gives rise to cell polarization and subsequent locomotion.

Towards the end of a Rho GTPase relaxation cycle, the membrane-bound active concentrations 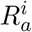 and 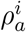 approach a uniform steady state over all vertices, terminating the peripheral gradients in 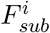 and thus the cell polarity (Fig. 2 A, B, *t/T* = 0.24 − 0.78 ; Fig. S3). The polarity ceases with the aid of increasing vertex angle magnitudes |*θ*^*i*^ | at e.g. the vertices *A* and *C* as follows: A strong protrusion at vertex *A* rapidly raises the activation rate 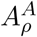 (Eq. S2) and in turn 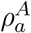 (Fig. 2 C), which suppresses protrusion by increasing the myosin motor number 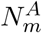 and thus the retrograde flow speed 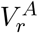 (Fig. 2 A, *t/T* = 0.24). Similarly, a strong contraction at vertex *C* increases the activation rate 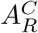 (Eq. S1) and surges 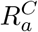 (Fig.2 D), which reverses the contraction (Fig.2 A, *t/T* = 0.24 − 0.78). Consequently, the two-way chemo-mechanical coupling in our model not only captures successive cell polarization and apolarization events but also replicates the complex cell morphology (Movies S1, S2).

### Whole-cell migration speed is governed by both substrate stiffness and viscosity

At the cell front, the radial spreading speed of a site 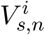 must be controlled by the substrate force *F*_*sub*_ upon molecular clutch formation with a time scale *τ*_*on*_ ≡ 1*/k*_*on*_. This is because *F*_*sub*_ sets the actin retrograde flow speed 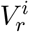 (Eq. 3), in turn determining 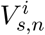 (Eq. 1) as a function of the stiffness *K*_*sub*_ and viscosity *γ*_*sub*_ of a uniform substrate, whereas the actin polymerization speed 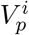 must remain largely unchanged. Then, an increase in *F*_*sub*_ increases 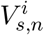, i.e., enhances traction on the substrate. For single FA dynamics, the role of the protrusion resistance force *F*_*p*_ in Eq. 2, 3 can be ignored since it builds up at a much slower rate than *F*_*sub*_ after clutch formation. Defining an elastic deformation time scale 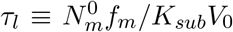 and a retardation time scale *τ*_*r*_ ≡ *γ*_*sub*_*/K*_*sub*_, the force generated by the myosin motors on the fully bound clutches (*P* ^*i*^ ∼ 1) sets the force on a uniform compliant substrate 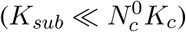 as

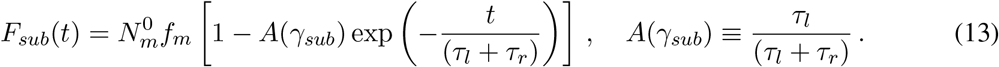

Eq. 13 is a single-motor-clutch relation derived from Eqs. 2 −4, 6 for 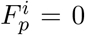 and holds until the time of force-dependent clutch dissociation *τ*_*off*_, i.e., at 0 ≤ *t* ≤ *τ*_*off*_ (Eq. 5, Sec. S4). Evidently, the time scales *τ*_*on*_, *τ*_*off*_, *τ*_*l*_, *τ*_*r*_ predominantly govern the focal adhesion dynamics and thus the migration speed.

We utilize our whole-cell model to explore the dependence of the average cell migration speed on the elastic stiffness *K*_*sub*_ and viscosity *γ*_*sub*_ of a uniform substrate. We interpret the simulation results with the help of Eq. 13. In our simulations, the nonuniform random initial conditions of the active Rac1 and RhoA concentrations 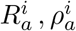 are the same as in the previous section (Sec. S3). Over one simulation with *M* ≫1 time steps, we define the average migration speed *V* as the nucleus trajectory length *L*≡ | **x**_*nuc*_(*M*Δ*t*) −**x**_*nuc*_(0) | divided by the total time *t*_*total*_ ≡ *M*Δ*t*, i.e., *V*≡ *L/t*_*total*_. We run *n* simulations to compute the distribution of *V* ; we define the sample mean migration speed as 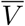.

For a uniform elastic substrate (*γ*_*sub*_ = 0), the speed-stiffness relation has been reported to show a biphasic profile [6, 7, 25, 40]. Our whole-cell simulations quantitatively reproduce the experimental dependence of 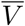 on *K*_*sub*_ (*n* = 5 ; Fig. 3 A) [7], which we explain by the competition between the two time scales *τ*_*l*_ and *τ*_*on*_ in Eq. 13 in the elastic limit (*τ*_*r*_ = 0). Importantly, the elastic deformation time scale *τ*_*l*_ can be interpreted as the characteristic clutch lifetime until the clutches break at *t* ≈ *τ*_*off*_ at a critical *F*_*sub*_, i.e., *τ*_*l*_ ≈ *τ*_*off*_ (Fig. S5 B, C). Because an FA site must generate optimal substrate force while the clutches are bound (*τ*_*on*_ ≲ *τ*_*l*_), the equality *τ*_*on*_ = *τ*_*l*_ defines a threshold stiffness 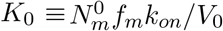, which allows us to classify a substrate as “stiff” when *K*_*sub*_ *> K*_0_ and “soft” when *K*_*sub*_ *< K*_0_ (Sec. S4). Based on the parameters in Table S2, *K*_0_ ≈ 8 *pN/nm*. On the stiff substrate (*K*_*sub*_ *> K*_0_, equivalently *τ*_*on*_ *> τ*_*l*_), because the bound clutch fraction is low (*P* ^*i*^ ≪1), the substrate traction force and thus the migration speed remain low (Fig. 3 A). On the soft substrate (*K*_*sub*_ *< K*_0_, equivalently *τ*_*on*_ *< τ*_*l*_), higher bound clutch fraction (*P* ^*i*^ ≈ 1) induces a higher traction force, and thus a higher migration speed (Fig. 3 A and B). On the very soft substrate (*K*_*sub*_ ≪ *K*_0_), Eq. 13 indicates that the traction force builds up very slowly at a time *τ*_*l*_ ≫*τ*_*on*_, suppressing the migration speed (Fig. 3 A, B). Next, we computed the average migration speed 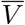 as a function of *K*_*sub*_ and *γ*_*sub*_ to resolve their combined effect on migration efficiency (Fig. 3 C–E). For lower viscosities (*γ*_*sub*_ = 0.1, 0.5 *pN ·s/nm*), the average speed preserves its biphasic profile about *K*_*sub*_ ≈ *K*_0_ (Fig. 3 C, D). At higher viscosities (*γ*_*sub*_ = 0.9 *pN s/nm*), however, 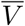 decreases monotonically with increasing stiffness (Fig. 3 C, D). Furthermore, 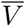 as a function of *γ*_*sub*_ is biphasic for a fixed stiffness *K*_*sub*_ *< K*_0_ (Fig. 3 D). The negligible change of the threshold stiffness *K*_0_ at lower viscosities and the non-monotonic dependence of 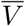 on *γ*_*sub*_ for *K*_*sub*_ *< K*_0_ can together be understood as follows: With increasing viscosity, whereas the viscoelastic deformation time scale *τ*_*l*_ + *τ*_*r*_ increases linearly, the force-dependent clutch lifetime *τ*_*off*_ changes only marginally from the purely elastic case (Fig. 3 F, S5 D). Therefore, the threshold stiffness *K*_0_ set by *τ*_*on*_ = *τ*_*l*_ remains nearly constant for sufficiently low viscosities (Fig. 3 C). Noting that the clutches in the fully bound state (*P* ^*i*^ ≈ 1) dissociate at a critical dimensionless force per clutch 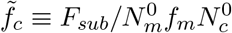 (≈ 7 ×10^−3^ for the parameters in Table S2; see Fig. 3 F), the viscosity can enhance the substrate traction only up until a threshold viscosity *γ*_0_ where, upon formation, each clutch can sustain a force 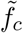 at most. Above *γ*_0_, because the substrate force per bond rapidly grows beyond 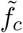 at a time shorter than *τ*_*on*_, the bond formation must be suppressed (*τ*_*off*_ *< τ*_*on*_), in turn leading to a significant traction loss (Fig. S5 D). Our numerical computation of the threshold viscosity *γ*_0_(*K*_*sub*_) based on this rationale agrees with the region of maximum migration speed in the contour plot (Fig. 3 C, full curve). Along the stiffness axis, *γ*_0_(*K*_*sub*_) is terminated at *K*_*sub*_ = *K*_0_, beyond which *τ*_*off*_ *< τ*_*on*_ independent of the viscosity of stiff substrates (Fig. 3 C, dashed line). This vertex-level scaling analysis explains the simulated whole-cell behavior in Fig. 3 C since the nonuniform initial 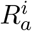 distribution breaks the cell symmetry and thus keeps 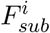 higher at the cell front than at the rear. Importantly, this difference in 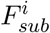 across the cell suppresses 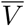 to be much lower than the vertex-level speed scale *V*_0_, enabling our whole-cell model to quantitatively capture the experimentally observed migration dynamics (see Fig. 3 A).

**Fig. 3:**
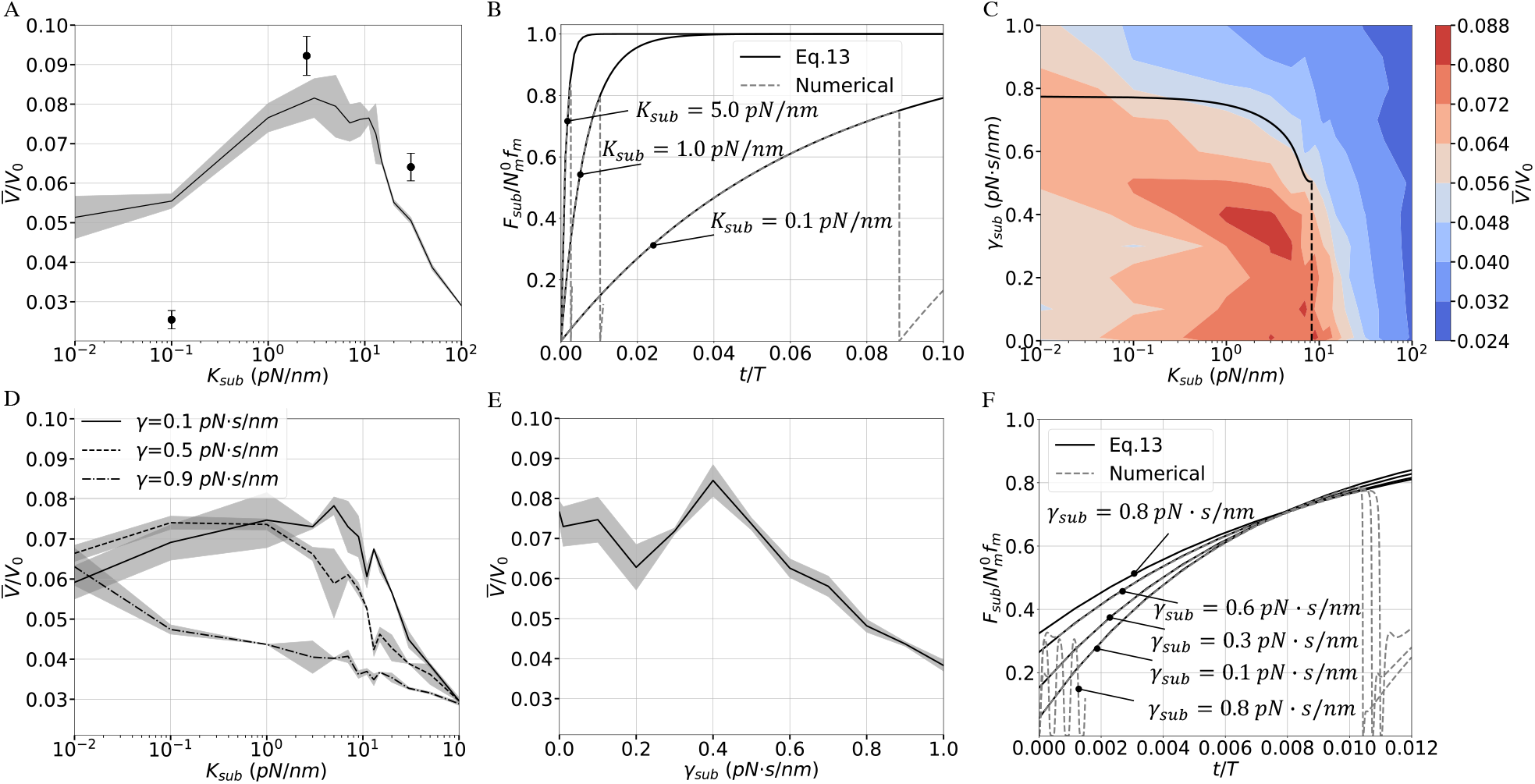
Cell migration speed on uniform viscoelastic substrates. (A) The dimensionless mean migration speed 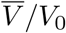 (full line) and its standard deviation (shaded area) as a function of the substrate elastic stiffness *K*_*sub*_ (the horizontal cross section of panel C at *γ*_*sub*_ = 0). For comparison, the dots and error bars are the experimental data from U373-MG human glioma cells on polyacrylamide hydrogels [7] [51]. (B) Analytical versus numerical dimensionless substrate force dynamics on an elastic substrate (*γ*_*sub*_ = 0). (C) Contour plot of 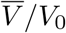 as a function of *K*_*sub*_ and substrate viscosity *γ*_*sub*_. The full curve is the threshold viscosity *γ*_0_, which is terminated at the threshold stiffness *K*_0_≈8*pN/nm* (dashed line). We ran *n* = 5 simulations to compute 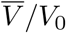 at each of the 180 points in the contour plot. (D) 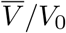 and its standard deviation (shaded areas) as a function of *K*_*sub*_ for finite *γ*_*sub*_. (E) 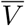 nd its standard deviation (shaded area) as a function of *γ*_*sub*_ for *K*_*sub*_ = 1 *pN/nm*. In panels A, D, E, the standard deviations are calculated over *n* = 5 simulations at each data point. (F) Analytical versus numerical dimensionless substrate force dynamics on a viscoelastic substrate (*K*_*sub*_ = 1.0 *pN/nm*). In panels B, F, the numerical profiles are obtained from an isolated sub-cellular motor-clutch model (Fig. S5).

### Migration direction depends on the local stiffness of nonuniform elastic substrates

Having elucidated how the substrate elastic stiffness and viscosity affect the migration speed, we next report cell symmetry breaking and directional migration on an elastic substrate (*γ*_*sub*_ = 0) with a stiffness gradient when chemically induced cell polarity is absent (Fig. 4 A). We specify the initial conditions for the membrane-bound Rho GTPase concentrations as follows: The active Rac1 concentration 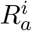 is assigned a random distribution, whereas the active RhoA concentration 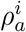 and the inactive Rac1, RhoA concentrations 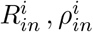 are set to be constant (Fig. S4). We characterize a nonuniform substrate by defining a reference stiffness *K*_*sub*,0_ at *x* = 0 where the nucleus is initially positioned (Fig. 4 A). We introduce a constant stiffness gradient ∇*K*_*sub*_ = 16 *pN/µm*^2^ in the +*x* direction; non-dimensionalization by the initial cell diameter 2*r*_0_ and the threshold stiffness *K*_0_ yields the unitless gradient 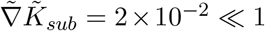.

**Fig. 4:**
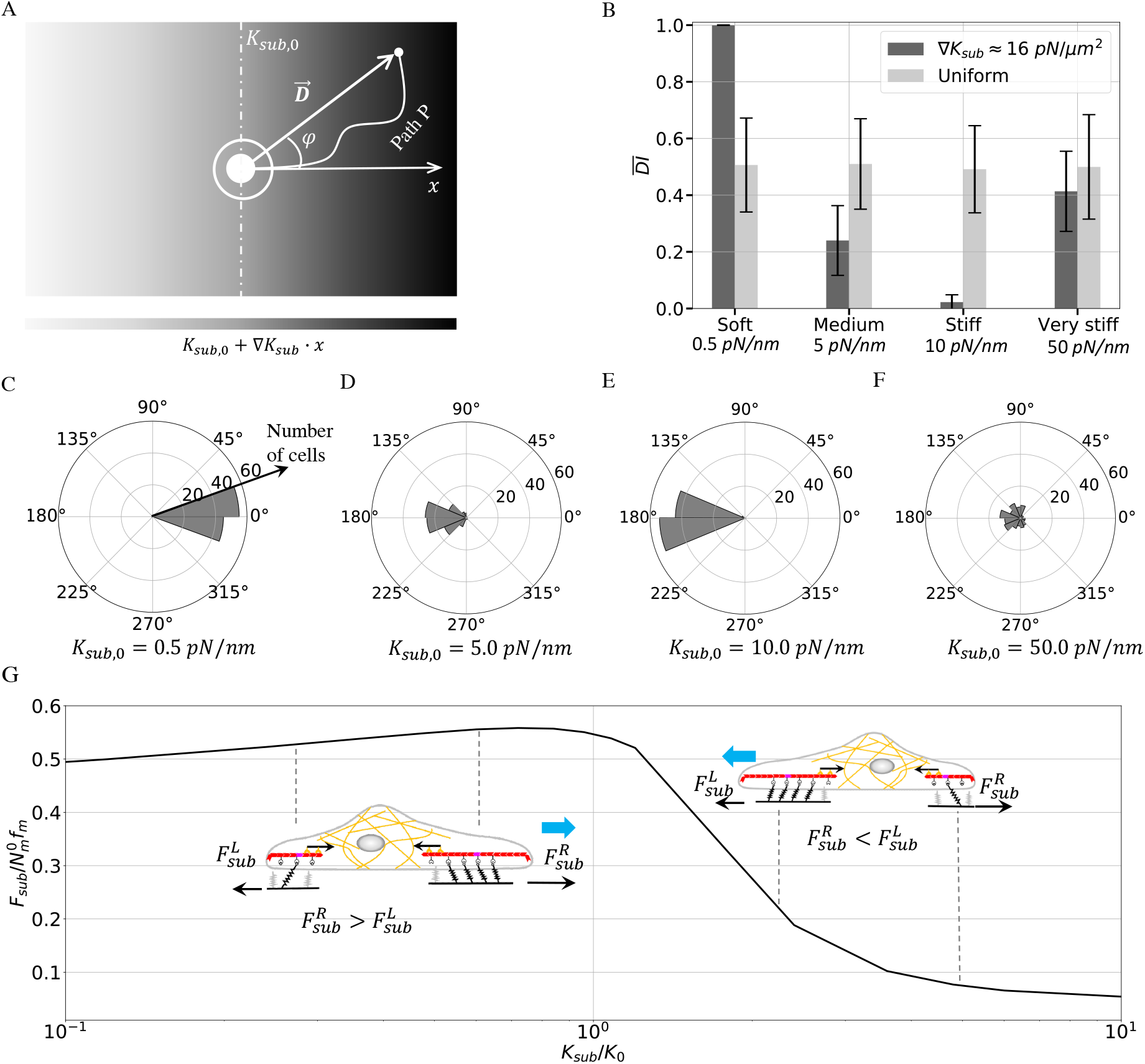
Migration direction on elastic substrates with stiffness gradients. (A) Simulation domain for a chemically apolar cell on a nonuniform substrate with a reference stiffness *K*_*sub*,0_ at *x* = 0 and constant stiffness gradient ∇*K*_*sub*_. The path *P* represents a migration event with a final distance 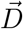 the origin and angular displacement *φ* with respect to the *x*-axis. (B) The mean durotactic index 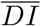 over *n* = 100 simulations for each of soft (*K*_*sub*,0_ *< K*_0_), medium (*K*_*sub*,0_ ≈*K*_0_), stiff (*K*_*sub*,0_ *> K*_0_), and very stiff (*K*_*sub*,0_ ≫*K*_0_) substrates. Light and dark bars represent 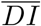 on uniform and nonuniform substrates with ∇*K*_*sub*_ = 16*pN/µm*^2^, respectively. Error bars show the standard deviations over *n* = 100 simulations. (C)–(F) depict the angular displacement (*φ*) distributions for each of the dark bars in B. The magnitude of each sector (with a 22.5^°^ bin size) represents the number of simulations in which the cell moved in the corresponding direction. (G) The biphasic local stiffness/substrate force relation from a vertex-level single motor-clutch model. The substrate traction forces at the leftmost and rightmost cell ends are defined as 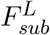 and 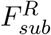. On nonuniform substrates, our simulations record three distinct migration behaviors, which we explain by comparing 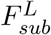 with 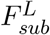 ; see the main text.

For soft (*K*_*sub*,0_ *< K*_0_), medium (*K*_*sub*,0_≈ *K*_0_), stiff (*K*_*sub*,0_ *> K*_0_), and very stiff (*K*_*sub*,0_ ≫ *K*_0_) substrates, we compare the effect of this small gradient on cell migration with that on a uniform substrate (*K*_*sub*_ = *K*_*sub*,0_). For each substrate configuration, we simulate *n* = 100 migration events. This allows us to statistically differentiate between durotaxis (migration up the stiffness gradient), anti-durotaxis (migration down the stiffness gradient), or adurotaxis (random motion irrespective of the stiffness gradient) by calculating the distributions of the durotactic index *DI* and the angular displacement *φ* (Fig. 4 A). We define *DI* as the net horizontal displacement over *Q* Rho-GTPase relaxation cycles in the gradient (+*x*) direction divided by the total horizontal displacement [15, 41], that is (ℋ: Heaviside step function, Δ*x*_*j*_ : displacement in the +*x* direction in the *j*^*th*^ cycle),

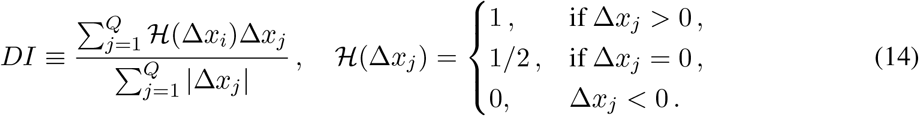

Eq. 14 returns *DI >* 0.5 for durotaxis, *DI* ≈0.5 for adurotaxis, and *DI <* 0.5 for anti-durotaxis. We compute the *DI* and *φ* distributions with *n* = 100 data points. We define the sample mean durotactic index as 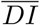 and the sample mean angular displacement as 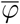.

Our results indicate that 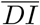 is inversely proportional to the reference substrate stiffness *K*_*sub*,0_ in the presence of a gradient. For ∇*K*_*sub*_ = 16 *pN/µm*^2^, the simulated cells undergo durotaxis on soft substrates, with 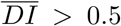 and 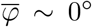 (Fig. 4 B, C), in agreement with experiments [16]. Higher gradients ∇*K*_*sub*_ = 32 *pN/µm*^2^ and ∇*K*_*sub*_ = 160 *pN/µm*^2^ (equivalently, 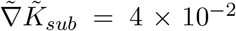 and 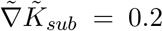 in dimensionless units) on soft substrates reinforce durotaxis, as evidenced by a robust increase in 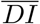 nd narrower angular distributions (Fig. S6). However, cells on medium and stiff substrates undergo anti-durotaxis with 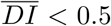, supported by the corresponding angular distributions (Fig. 4 B, D, E). These results agree well with the stiffness-sensitive migration behaviors of the U-251MG glioblastoma and talin-low MDA-MB-231 breast cancer cells [18]. On very stiff nonuniform substrates (*K*_*sub*,0_ ≫*K*_0_), the cells undergo adurotaxis (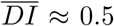: uniform), i.e., follow random migration trajectories (Fig. 4 B, F). Our simulations for substrates with uniform stiffness also yield 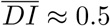 ;the absence of cell symmetry-breaking again leads to adurotaxis (Fig. 4 B).

The capability of our whole-cell model to produce durotaxis, anti-durotaxis, and adurotaxis as a function of the substrate stiffness and its gradients allows us to propose a unifying mechanism for stiffness-sensitive migration. On one hand, on a soft nonuniform substrate, an FA site attached to a stiffer region generates higher traction force where the cell in turn spreads faster than at another site attached to a softer region. In other words, 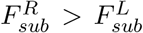 drives 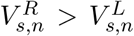 in Fig. 4 G, navigating the cell up the stiffness gradient (durotaxis). On the other hand, on a stiff nonuniform substrate, the cell generates a higher traction force on the softer region 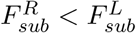 drives 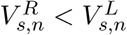, migrating down the stiffness gradient (anti-durotaxis). Adurotaxis occurs when the cell becomes insensitive to the gradients on very stiff substrates or when the traction forces are balanced at the opposing sides of the cell on a substrate with *K*_*sub*,0_ ≲ *K*_0_ (Fig. 4 G). Therefore, the biphasic dependence of the substrate force on the local stiffness is sufficient for the emergence of different durotactic phenotypes on nonuniform substrates.

### Migration direction depends on both viscosity and stiffness of viscoelastic substrates

We have reported that the substrate viscosity *γ*_*sub*_ alters the non-monotonic dependence of the migration speed on the substrate stiffness *K*_*sub*_. Here, we study how uniform *γ*_*sub*_ alters the directional migration of a chemically apolar cell in the presence of a constant stiffness gradient ∇*K*_*sub*_ = 16*pN/µm*^2^ in the +*x* direction. The initial conditions are the same as in the preceding section. To plot the contour map in Fig. 5 A, we performed *n* = 5 simulations to obtain the mean durotactic index 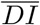 for each data point *K*_*sub*,0_, *γ*_*sub*_ that specifies a viscoelastic substrate. Fig. 5 A exhibits three distinct migration regimes as a result of the mechanical interactions between the cell and the medium: Regime I pervades soft substrates (*K*_*sub*,0_ *< K*_0_) with a sufficiently low viscosity where the cell undergoes durotaxis (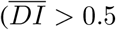 and 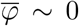 ; Fig. 5 A–D), in accordance with Fig. 4 C. Increasing *K*_*sub*,0_ or *γ*_*sub*_ leads to Regime II in which the cell sustains anti-durotaxis (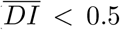 and 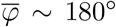 Fig. 5 A–C, E). This is because the magnitude and duration of the substrate traction forces at the FA sites are reduced on stiff and more viscous substrates concurrently with shorter clutch lifetimes, directing the cell towards softer regions (see Fig. 3 F and Fig. 4 G). Because cells are insensitive to a very high stiffness (*K*_*sub*,0_ ≫*K*_0_) or viscosity (*γ*_*sub*_ *> γ*_0_), they exhibit adurotaxis in this limit, i.e., move randomly with small displacements, which we identify as Regime III (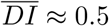 and φ: uniform; Fig. 5 A–C, F). Overall, Fig. 5 A generalizes our earlier findings in Fig. 3 C for the mesencyhmal migration on uniform viscoelastic substrates.

**Fig. 5:**
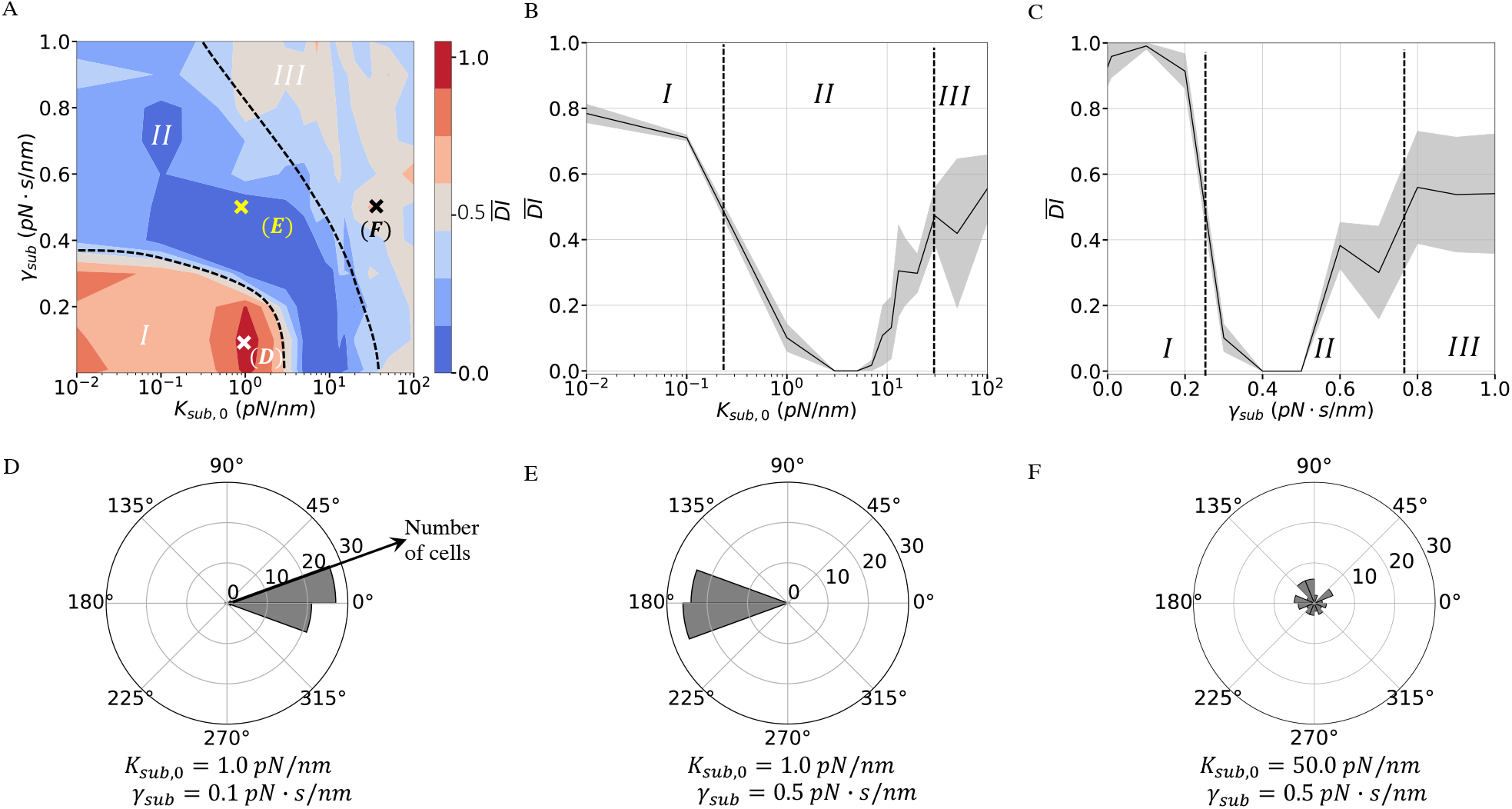
Migration direction on viscoelastic substrates with stiffness gradients. (A) Contour plot of the mean durotactic index 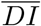 as a function of the reference substrate stiffness *K*_*sub*,0_ and uniform substrate viscosity *γ*_*sub*_. The contour plot contains 180 data points with *n* = 5 at each point. As guides to the eye, the dashed black curves divide the contour plot into three migration regimes: Regime I (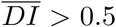 ; durotaxis), Regime II (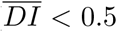; anti-durotaxis), Regime III (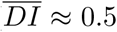 ; adurotaxis). (B) The mean durotactic index 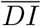versus the reference stiffness *K*_*sub*,0_ with a constant substrate viscosity *γ*_*sub*_ = 0.3 *pN s/nm*. (C) The mean durotactic index 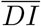 versus the viscosity *γ*_*sub*_ when *K*_*sub*,0_ = 1.0 *pN/nm*. Shaded regions in B and C represent the standard deviations for *n* = 5. (D)–(F) depict the angular displacement (*φ*) distributions at each cross symbol in A. Each sector with a 22.5^°^ bin size represents the number of simulations in which the cell moved in the corresponding direction for *n* = 50.

### RhoA induced actomyosin contractility alters cell migration direction

As opposed to our analysis of a chemically apolar cell migrating on a nonuniform viscoelastic substrate (Fig. 5), directional migration in biological environments is regulated by both mechanical and chemical cues [42, 43]. To address this complexity, we consider that external chemical fields can trigger a biased active RhoA distribution 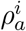 at the cell membrane [24, 42]. Taking a nonuniform distribution of 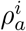 as an initial condition instantaneously generates a localized actomyosin contractility due to the high retrograde flow (Fig. 6 A, B). On a soft elastic substrate with *K*_*sub*,0_ = 1*pN/nm < K*_0_ and ∇*K*_*sub*_ = 16*pN/µm*^2^, the simulated cells therefore exhibit anti-durotaxis (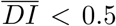 and 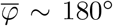 for *n* = 50) in contrast with the durotactic migration in the absence of RhoA induced contractility (Fig. 6 C, D). Further simulations on viscoelastic substrates demonstrate that the chemical polarity can override the cell-matrix mechanical interactions, making cells migrate against the stiffness gradient regardless of the viscosity magnitudes (Fig. 6 E–H versus Fig. 5 D–F). These results suggest a selection mechanism for the migration direction in the presence of both the mechanical and chemical fields *in vivo*.

**Fig. 6:**
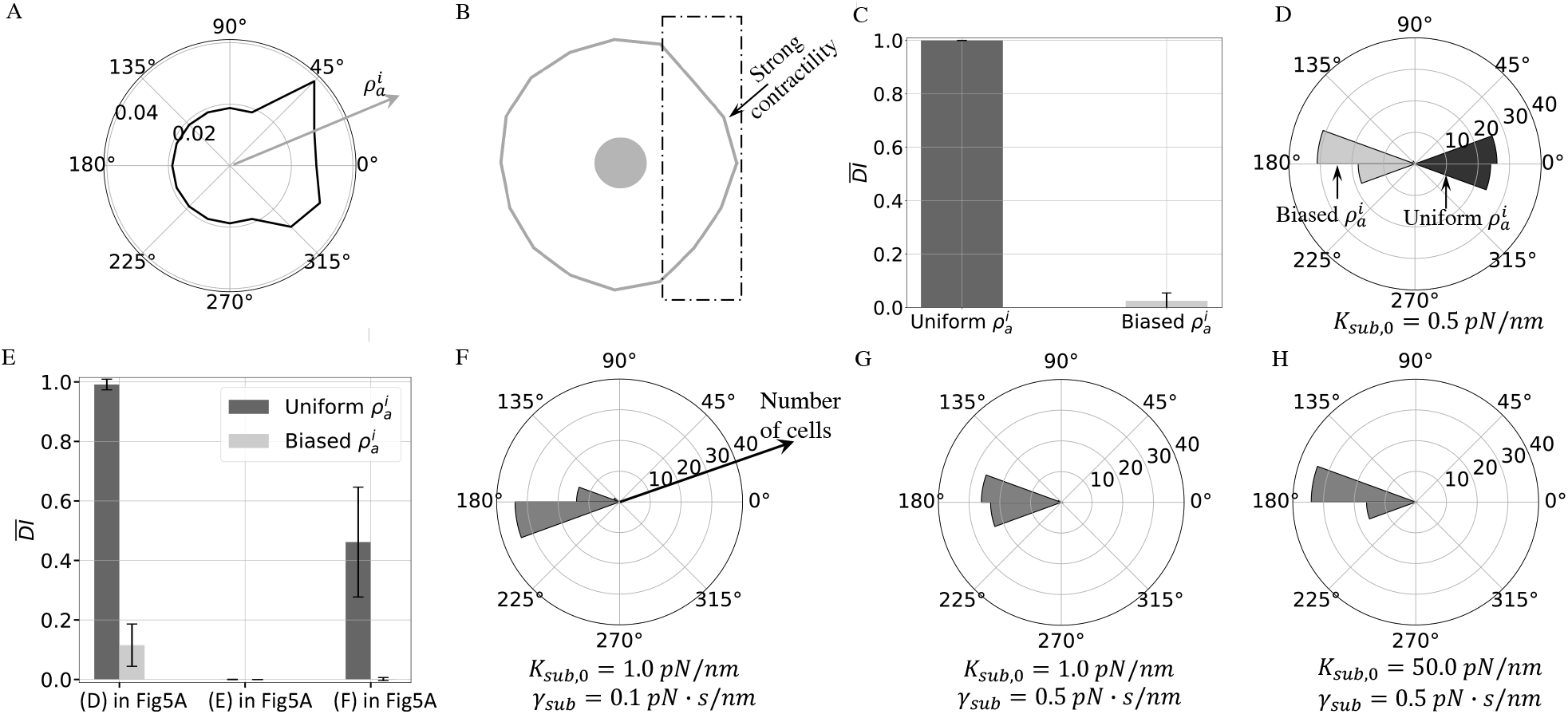
Rho GTPase signals can reverse the mechanically favored migration direction. (A) Initialcondition of the active RhoA concentration 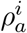 on the cell vertices. (B) The resulting actomyosin induced cell contraction. (C) The mean durotactic index 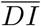 of cells with the uniform RhoA and biased RhoA signals on the nonuniform elastic substrates. Error bars are the standard deviations over *n* = 50 simulations each. (D) The angular displacement (*φ*) distributions with the uniform and biased 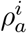, illustrated by black and gray sectors, respectively (*n* = 50 simulations each). Each sector with a 22.5^°^ bin size represents the number of simulations where the cell moved in the corresponding direction. (E) On viscoelastic substrates specified by the cross symbols in Fig. 5 A, 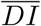 for *n* = 50. Light and dark bars represent 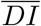 of uniform and biased RhoA signals 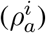, respectively. Error bars show the standard deviations for *n* = 50. (F)–(H) depict the angular displacement (*φ*) distributions for each of the light bars in E.

## Discussion

Mesenchymal migration involves a dynamical interplay between physical and chemical processes across the sub-cellular and cellular scales in response to the biological viscoelastic media. To account for this complexity, our whole-cell model couples the dynamics of sub-cellular FA sites, which probe the local mechanical properties of the surrounding medium, with the local membrane mechanics and the cell-level reaction-diffusion dynamics of the two relevant Rho GTPase signals, Rac1 and RhoA. At the cellular scale of our model, the cytoskeleton transmits the forces between the FA sites and the cell nucleus where the global force balance sets the migration direction and speed (Fig. 1). That way, our whole-cell model exhibits all distinct stages of mesenchymal migration, i.e., the emergence of Rho GTPase induced cell polarity following FA formation, the associated complex shape changes, and the subsequent migration on uniform substrates via a stick-slip motion evidenced by the oscillating local substrate traction forces (Fig. 2). For uniform elastic substrates, we analytically determined a threshold stiffness *K*_0_, which classifies the substrates as soft (*K*_*sub*_ *< K*_0_), medium (*K*_*sub*_ ≈ *K*_0_), stiff (*K*_*sub*_ *> K*_0_), and very stiff (*K*_*sub*_ ≫ *K*_0_). The migration speed is non-monotonic with respect to the substrate stiffness about *K*_0_, in agreement with experiments. We further found that, for viscoelastic substrates, this biphasic trend is terminated at a threshold viscosity *γ*_0_ where the rapid traction force build-up cannot be sustained by the bound molecular clutches anymore, suppressing their formation in the first place (Fig. 3). Next, we investigated the effect of the stiffness gradients on migration by switching off the Rho GTPase induced cell polarity. Our results demonstrate that the biphasic dependence of the local spreading speed on the local stiffness gives rise to durotaxis on soft substrates with low viscosity, anti-durotaxis on stiff or viscous substrates, or adurotaxis on very stiff substrates (Fig. 4, 5). Finally, we showed that imposing a RhoA-driven strong local contraction, which can be experimentally achieved by external chemical cues, can steer the cell along the Rho GTPase induced polarity irrespective of the stiffness gradients (Fig. 6).

We close with a discussion of the limitations of our whole-cell model. Unlike the classical motor-clutch model that we used here for the FA dynamics, experiments have found an adhesion reinforcement effect due to talin unfolding at the clutches [44]. An augmented clutch model that considers the adhesion reinforcement has replicated the monotonic relation between the substrate traction force and substrate stiffness as well as that between the cell area and stiffness [13, 26, 45]. However, this theoretical modification cannot preserve the experimentally confirmed biphasic speed versus stiffness profile, which is a natural theoretical question for further investigation.

Another limitation is that we only track the activities of two primary chemical signals, Rac1 and RhoA, for simplicity. However, Rac1 and RhoA interact with other proteins such as ROCK and DIA [22]. Moreover, we manually reinitialize the active Rac1 and RhoA concentrations when they reach a steady state. A self-perpetuating oscillatory state in signal transduction can be obtained by taking into account regulations of other proteins [22], inputs from ECM [46], and a more comprehensive reverse coupling from the cell mechanics [30]. Although this simplification does not undermine our results, integrating the dynamics of additional proteins can enhance the chemo-mechanical feedback of our whole-cell model.

A third shortcoming is that our whole-cell model simplifies the cell structural mechanics. We represent the cytoskeleton by linear Hookean springs. However, since the cell stiffness depends on the cell area [45], large deformations may induce a nonlinear strain-stiffening effect of the cytoskeleton network. It is also known that cytoskeletal filaments are inherently dynamic due to their polymerization and crosslinking kinetics [47]. Furthermore, whereas our model assumes static links between the nucleus and cytoskeleton, the nucleus motion is affected by constant cytoskeletal–nuclear binding and unbinding events [48]. Incorporating the cytoskeletal kinetics and cytoskeletal-nuclear interactions into our whole-cell model may allow us to assess their relevance to the cell migration dynamics.

These limitations notwithstanding, our whole-cell model unifies seemingly opposing durotactic behaviors due to the mechanical cell-medium interactions and the intracellular signaling pathways. It therefore elucidates complex mechano-sensing at the single-cell level and paves the way for the generalization of cell behavior and migration in three dimensional microenvironments.

## Author contributions

W.S.: conceptualization, formal analysis, investigation, software, validation, visualization, writing – original draft, review and editing; C.N.K.: conceptualization, formal analysis, project administration, resources, validation, writing – original draft, review and editing.

## Data availability

The source codes of the simulations can be downloaded at https://github.com/_nadirkaplan/migration_multiscale_theory.

## Acknowledgments

We thank the College of Science at Virginia Tech for financial support.

