## Supporting Information for "A multiscale whole-cell theory for mechano-sensitive migration on viscoelastic substrates"

Wenya Shu[1,2] and C. Nadir Kaplan[1,2 *]
[1]Department of Physics,
Virginia Polytechnic Institute and State University, Blacksburg, VA 24061, USA.
[2]Center for Soft Matter and Biological Physics,
Virginia Polytechnic Institute and State University, Blacksburg, VA 24061, USA.

#### Supplementary Movies

**Movie S1 - Cell polarity and migration on a soft substrate.** Chemically induced cell polarization and migration with a complex morphology on a uniform elastic substrate with a stiffness $K_{sub} = 3\,pN/nm$). The initial conditions for the Rho GTPase protein concentrations are given in Fig. S3.

**Movie S2 - Cell polarity and migration on a stiff substrate.** Chemically induced cell polarization and migration with a complex morphology on a uniform elastic substrate with a stiffness $K_{sub} = 100\,pN/nm$). The initial conditions for the Rho GTPase protein concentrations are given in Fig. S3.

#### S1 Chemo-mechanical coefficients governing Rho GTPase dynamics

The reaction-diffusion model of the Rho GTPase dynamics is based on the well-acknowledged mutual inhibition and autoactivation effects in Rac-Rho interactions. Additionally, feedback between cell mechanical forces and signaling pathways is essential for the cell motility and its oscillatory shape changes. In this study, definitions of rates $A_G^i$ in Eq. 12 accommodate autoactivation and antagonistic effects as well as feedback from the mechanical deformations of the cell membrane. The Rac1 activation rate $A_R^i$ at the $i^{th}$ vertex is composed of three terms,

$$A_R^i = \frac{\alpha_R {R_a^i}^3}{\left({R_0}^3 + {R_a^i}^3\right)} + \frac{\beta_R {\rho_0}^3}{\left({\rho_0}^3 + {\rho_a^i}^3\right)} + e^{\mathcal{H}(\Delta\theta - c_0\theta_0)\frac{\Delta\theta}{\theta_0}} K_b^+ \,, \tag{S1}$$

where $R_0$ and $\rho_0$ are reference levels of the active Rac1 and RhoA on vertex, respectively (Table S2). The first term takes into account positive feedback from the active Rac1 to itself. The magnitude of the autoactivation effect is represented by $\alpha_R$ [1, 2]. The second term describes the mutual inhibition effect between Rac1 and RhoA with a rate $\beta_R$ [1, 2]. The last term couples the active level of Rac1 to mechanical deformations of the cell at the $i^{th}$ vertex. $\mathcal{H}()$ denotes the Heaviside step function. The initial angle at a cell vertex is labeled by $\theta$ , and $\Delta\theta \equiv \theta - \theta_0$ denotes the change in angle at the vertex. Here we assume that the active rate of Rac1 soars once the increase of the vertex angle exceeds the threshold $c_0\theta_0$ , due to the cell contraction. Similarly, RhoA activation rate $A_\rho^i$ at the $i^{th}$ vertex also consists of three terms,

$$A_\rho^i = \frac{\alpha_\rho {\rho_a^i}^3}{\left({\rho_0}^3 + {\rho_a^i}^3\right)} + \frac{\beta_\rho {R_0}^3}{\left({R_0}^3 + {R_a^i}^3\right)} + e^{\mathcal{H}(\Delta\theta + c_0\theta_0)\frac{\Delta\theta}{\theta_0}} \kappa_b^+ . \tag{S2}$$

We assume that the active rate of RhoA increases once the reduction of the vertex angle satisfies $\Delta\theta_0 < -c_0\theta_0$) due to the cell polarization. The numerical values of all the parameters in Eq S1, S2 are listed in Table S2. Coupling Rac1 and RhoA to mechanical deformations enables recovery of cell deformations and recurrent polarization.

### S2 Mesh refinement study

All equations are non-dimensionalized with the scaling factors that are chosen as $X = 2\pi r_0$, $T = X/V$, $F = N_m^0 f_m$ and $V = V_0$. The primary equations in both dimensional and dimensionless forms are given in Table. S1. The values of constant parameters are given in Table.S2. Implementation of the proposed model is schematically illustrated in the flowchart in Fig. S1.

A typical problem of the vertex model is the strong dependence of the cell migration dynamics on the mesh refinement level (i.e., the number of vertices $N$). A well-established model should give a unique and convergent result with increasing mesh refinement. Thus, the mesh refinement study is essential to determine the adequate number of mesh points. We consider several mesh refinement levels with the number of vertices being the multiples of 8. As a reference, we assume that there are 16 focal adhesion (FA) sites at the cell periphery. Based on this assumption, the cytoskeleton stiffness $K_{ck}$ is multiplied by $16/N$ to ensure that the level of membrane discretization does not change the cytoskeletal elastic energy of the deformed cell. The membrane spring stiffness ($K_m$) is inversely proportional to the spring length between adjacent mesh points and is thus multiplied by $N/16$. Additionally, the FA size is scaled by $16/N$ . Correspondingly, the substrate stiffness ($K_{sub}$) , viscosity ($\gamma_{sub}$), the number of the clutches $N_c$ and the number of myosin motors $N_m$ should also be multiplied by $16/N$. The parameters of the model with $N = 16$ nodes are given in Table.S2.

Chemically induced cell symmetry breaking is the result of the initially polarized Rac1 signal on the membrane with the non-uniform distribution in Fig. S2A. Once the inactive level of Rac1 protein in the cytosol settles to the steady state, the Rac1 signal is initialized to the state at $t = 0$. The resulting dynamics in simulations yield periodic and small amplitude fluctuations of the cell area and persistent migration. The elastic energy for the membrane and cytoskeleton system is given by

$$\Psi = \sum_{i=1}^{N-1} \frac{1}{2} K_m \left(l_+^i - l_0\right)^2 + \sum_{i=1}^{N} \frac{1}{2} K_{cs} \left(r^i - r_0\right)^2 . \tag{S3}$$

We plotted the area, average migration velocity, and the elastic energy with respect to the mesh size N in Fig. S2B-D. The model provides a unique and convergent solution of the cell migration with the increasing mesh size. In particular, the model with $N = 16 - 24$ vertices is adequate to ensure accuracy and also achieve fair computational efficiency in the simulations.

Table S1: Governing equations in dimensional and dimensionless forms.

Let $t = \tilde{t}T,\ x_i = \tilde{x}_i X,\ f = \tilde{f}F$ where $X = 2\pi r_0, T = \frac{X}{V_0}, F = N_m^0 f_m$.

| Eq # | Dimensional | Dimensionless |
|---|---|---|
| $\tilde{t} = \frac{t}{T},\ \tilde{k}_{on} = k_{on}T, \tilde{\alpha} = \alpha F T,\ \tilde{k}_s = k_s T,\ \tilde{k}_c = k_c T, \tilde{f}_c = \frac{f_c}{F},\ \tilde{x} = \frac{x}{X}, \tilde{f} = \frac{f}{F},$ $\tilde{\gamma} = \frac{\gamma}{FT/X}, \tilde{K}_{sub} = \frac{K_{sub}}{F/X}, \tilde{K}_c = \frac{K_c}{F/X},\ \tilde{V} = \frac{V}{V_0}$ | | |
| (2-3) | $\dot{x}_r^i = V_0\left(1 - \frac{P^i N_c^i K_c\left(x_r^i - x_{sub}^i\right) - F_p^i}{N_m^i f_m}\right)$ | $\dot{\tilde{x}}_r^i = \left(1 - \frac{P^i N_c^i \tilde{K}_c\left(\tilde{x}_r^i - \tilde{x}_{sub}^i\right) - \tilde{F}_p^i}{N_m^i / N_m^0}\right)$ |
| (5) | $\frac{dP^i}{dt} = k_{on}^i\left(1 - P^i\right) - k_{off}^i P^i$ | $\frac{dP^i}{d\tilde{t}} = \tilde{k}_{on}^i\left(1 - P^i\right) - \tilde{k}_{off}^i P^i$ |
| (6) | $\gamma_{sub}^i \dot{x}_{sub}^i + K_{sub}^i x_{sub}^i = P^i N_c^i K_c\left(x_r^i - x_{sub}^i\right)$ | $\tilde{\gamma}_{sub}^i \dot{\tilde{x}}_{sub}^i + \tilde{K}_{sub}^i \tilde{x}_{sub}^i = P^i N_c^i \tilde{K}_c\left(\tilde{x}_r^i - \tilde{x}_{sub}^i\right)$ |
| $\tilde{K}_m = \frac{K_m}{F/X}, \tilde{K}_{cs} = \frac{K_{cs}}{F/X}, \tilde{\eta}_m^i = \frac{\eta_m^i V_0}{F/X},\ \tilde{\eta}_{cp} = \frac{\eta_{cp} V_0}{F/X}, \tilde{l}^i = \frac{l^i}{X}, \tilde{r} = \frac{r}{X}$ | | |
| (8-9) | $\begin{cases} F_P^i + F_{cs}^i + \boldsymbol{F}_{\boldsymbol{m}\pm}^i \cdot \boldsymbol{n}^i - \eta_m^i \langle l^i \rangle V_{s,n}^i = 0 \\ \boldsymbol{F}_{\boldsymbol{m}\pm}^i \cdot \boldsymbol{\tau}^i - \eta_m^i \langle l^i \rangle V_{s,\tau}^i = 0 \end{cases}$ | $\begin{cases} \tilde{F}_P^i + \tilde{F}_{cs}^i + \tilde{\boldsymbol{F}}_{\boldsymbol{m}\pm}^i \cdot \boldsymbol{n}^i - \tilde{\eta}_m^i \langle \tilde{l}^i \rangle \tilde{V}_{s,n}^i = 0 \\ \tilde{\boldsymbol{F}}_{\boldsymbol{m}\pm}^i \cdot \boldsymbol{\tau}^i - \tilde{\eta}_m^i \langle \tilde{l}^i \rangle \tilde{V}_{s,\tau}^i = 0 \end{cases}$ |
| (10-11) | $\sum_{i=1}^{N}\left(F_{am}^i + F_{cs}^i\right)\boldsymbol{n}^i + \boldsymbol{F}_{\boldsymbol{nuc}} = 0$ | $\sum_{i=1}^{N}\left(\tilde{F}_{am}^i + \tilde{F}_{cs}^i\right)\boldsymbol{n}^i - 6\pi\tilde{\eta}_{cp}\tilde{r}_{nuc}\tilde{\boldsymbol{V}}_{\boldsymbol{nuc}} = 0$ |
| $\tilde{D} = \frac{D}{XV_0},\ \tilde{\alpha}_G = \alpha_G T,\ \tilde{\beta}_G = \beta_G T,\ \tilde{K}_b^+ = K_b^+ T,\ \tilde{\kappa}_b^+ = \kappa_b^+ T,\ \tilde{I}_G^i = I_G^i T, \tilde{M}_G^+ = M_G^+ T, \tilde{M}_G^- = M_G^- T$ | | |
| (12) | $J_y^i = -D\left(\frac{G_y^{i+1}/\langle l^{i+1}\rangle - G_y^i/\langle l^i\rangle}{\lvert \mathbf{l}_+^i \rvert}\right)$ $\begin{cases} \frac{dG_a^i}{dt} = A_G^i G_{in}^i - I_G^i G_a^i + \left(J_a^{i-1} - J_a^i\right) \\ \frac{dG_{in}^i}{dt} = -A_G^i G_{in}^i + I_G^i G_a^i + \left(J_{in}^{i-1} - J_{in}^i\right) + \frac{M_G^+ G_{cp}}{N} - M_G^- G_{in}^i \\ \frac{dG_{cp}}{dt} = -M_G^+ G_{cp} + \sum_{i=1}^{N} M_G^- G_{in}^i \end{cases}$ | $\tilde{J}_y^i = -\tilde{D}\left(\frac{G_y^{i+1}/\langle \tilde{l}^{i+1}\rangle - G_y^i/\langle \tilde{l}^i\rangle}{\lvert \tilde{\mathbf{l}}_+^i \rvert}\right)$ $\begin{cases} \frac{dG_a^i}{d\tilde{t}} = \tilde{A}_G^i G_{in}^i - \tilde{I}_G^i G_a^i + \left(\tilde{J}_a^{i-1} - \tilde{J}_a^i\right) \\ \frac{dG_{in}^i}{d\tilde{t}} = -\tilde{A}_G^i G_{in}^i + \tilde{I}_G^i G_a^i + \left(\tilde{J}_{in}^{i-1} - \tilde{J}_{in}^i\right) + \frac{\tilde{M}_G^+ G_{cp}}{N} - \tilde{M}_G^- G_{in}^i \\ \frac{dG_{cp}}{d\tilde{t}} = -\tilde{M}_G^+ G_{cp} + \sum_{i=1}^{N} \tilde{M}_G^- G_{in}^i \end{cases}$ |

Table S2: Fixed model parameters used in the simulations.

| Symbol | Parameters | Dimensional | Dimensionless |
|---|---|---|---|
| $f_m$ | Single myosin motor stall force | 2.0 $pN$ [3, 4] | $1/100$ |
| $K_c$ | Clutch stiffness | 2.0 $pN/nm$ [3, 5] | $100\pi$ |
| $k_{on}^i$ | Rate constant of clutch association | 5.0 $s^{-1}$ [4] | $1250\pi/3$ |
| $k_{r0}$ | Clutch unloaded off-rate | 0.25 $s^{-1}$ [4, 5] | $125\pi/6$ |
| $k_{c0}$ | Clutch unloaded catch-rate | 120 $s^{-1}$ [4, 5] | $10000\pi$ |
| $f_{c0}$ | Characteristic catch force | 0.5 $pN$ [4] | $1/400$ |
| $f_{r0}$ | Characteristic rupture force | 1.0 $pN$ [4] | $1/200$ |
| $V_0$ | Unloaded retrograde flow velocity | 120.0 $nm/s$ [3] | 1.0 |
| $V_p^0$ | Characteristic polymerization rate | 120.0 $nm/s$ [6] | 1.0 |
| $r_0$ | Cell radius | 5.0 $\mu m$ [6] | $1/2\pi$ |
| $r_{nuc}$ | Nucleus radius | 2.0 $\mu m$ [7, 8] | $1/10\pi$ |
| $K_m$ | Membrane stiffness | 2.0 $pN/\mu m$ [9] | $\pi/10$ |
| $\eta_m^i$ | Viscosity of the surrounding medium | 10.0 $Pa \cdot s$ [10] | $3\pi/50$ |
| $K_{cs}$ | Cytoskeletal filament stiffness | 20.0 $pN/\mu m$ [11] | $\pi$ |
| $N_c^0$ | Reference molecular clutch number | 100 [4] | 100 |
| $N_m^0$ | Reference myosin motor number | 100 [4] | 100 |
| $M_R^+$, $M_\rho^+$ | Rac1, RhoA membrane association rate | 0.04 $s^{-1}$ [12] | $10\pi/3$ |
| $M_R^-$, $M_\rho^-$ | Rac1, RhoA membrane dissociation rate | 0.04 $s^{-1}$ [12] | $10\pi/3$ |
| $K_b^+$ | Baseline Rac1 activation rate | 0.6 $s^{-1}$ | $50\pi$ |
| $\kappa_b^+$ | Baseline RhoA activation rate | 0.6 $s^{-1}$ | $50\pi$ |
| $K^-$ | Rac1 deactivation rate ($I_R^i = K^-$) | 0.9 $s^{-1}$ [1, 13] | $75\pi$ |
| $\kappa^-$ | RhoA deactivation rate ($I_\rho^i = \kappa^-$) | 0.9 $s^{-1}$ [1, 13] | $75\pi$ |
| $R_0$ | Reference level of the active Rac1 | 3/160 | 3/160 |
| $\rho_0$ | Reference level of the active RhoA | 3/160 | 3/160 |
| $\theta_0$ | Vertex angle at $t = 0$ | $7\pi/8$ | $7\pi/8$ |
| $c_0$ | Ratio of the threshold angle | 0.3 | 0.3 |
| $\alpha_R$ | Positive feedback rate on Rac1 | 0.3 $s^{-1}$ [1] | $75\pi/3$ |
| $\alpha_\rho$ | Positive feedback rate on RhoA | 0.3 $s^{-1}$ [1] | $75\pi/3$ |
| $\beta_R$ | Rate of Rac1 inhibition by RhoA | 0.3 $s^{-1}$ [14] | $75\pi/3$ |
| $\beta_\rho$ | Rate of RhoA inhibition by Rac1 | 0.3 $s^{-1}$ [14] | $75\pi/3$ |
| $D$ | Diffusivity on the membrane | 0.01 $\mu m^2/s$ [1] | $1/120\pi$ |
| $\eta_{cp}$ | Cytoplasm viscosity | 800 $Pa \cdot s$ [15] | 4.8$\pi$ |

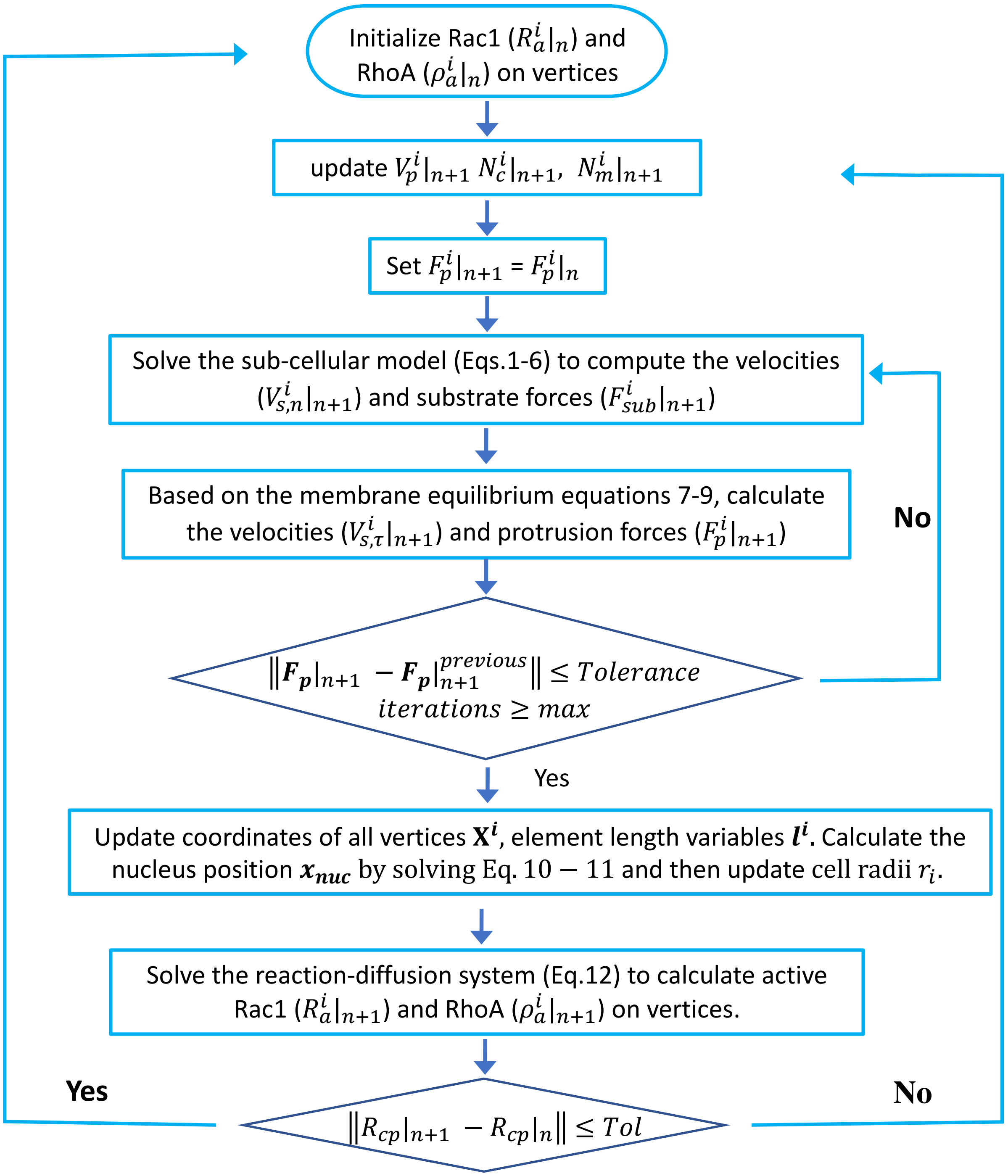


Fig. S1: Flowchart of the computational algorithm for the multiscale whole-cell model.

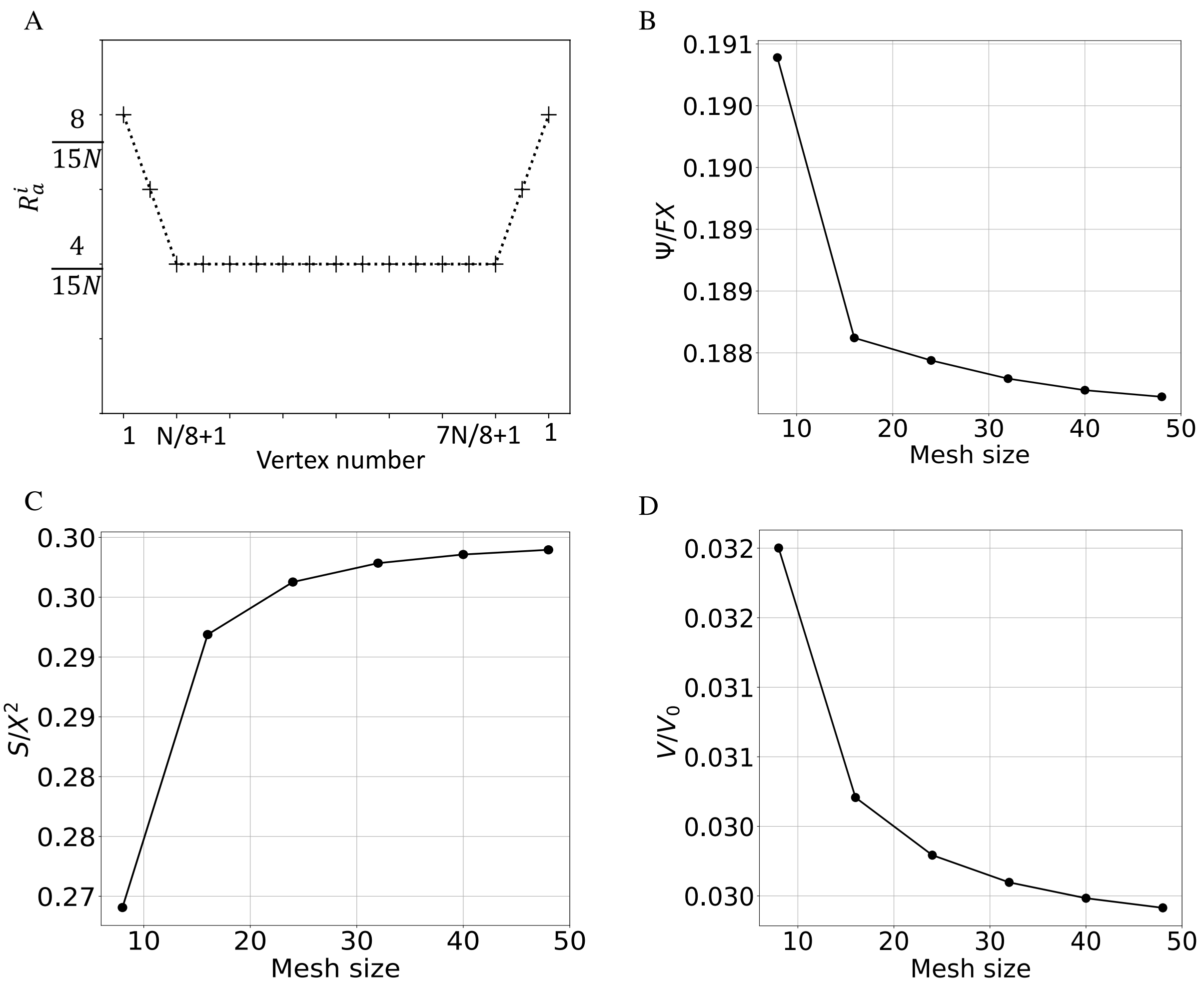


Fig. S2: **Mesh convergence study.** (A) Initial concentrations of the Rac1 proteins on vertices. (B) Mesh size dependence of the dimensionless elastic energy (Eq. S3). (C) Mesh size dependence of the dimensionless cell area $S/X$. (D) Mesh size dependence of dimensionless the migration speed.

### S3 Initial conditions of Rho GTPase protein concentrations

The Rho GTPase proteins are the central regulators of cell migration. Initial conditions of Rac1 and RhoA control both the morphology and migratory directionality of the cell. The following two initial conditions are specified to study cell mechano-sensitivity. To study the effect of ECM mechanics on the cell migration speed on uniform substrates, the polarized Rac1 signal with random noise was defined as the initial condition, i.e., 1-2 vertices among the rightmost five vertices have at least one-time stronger active Rac1 signal than the other vertices (see Fig.S3A). Other types of the membrane bound proteins ($\rho_a^i$, $R_{in}^i$, $\rho_{in}^i$) are randomly distributed on vertices (Fig.S3B-D). Since the total concentration of Rac1 and RhoA are each conserved, the initial cytoplasmic Rac1 concentration $R_{cp}(t=0)$ and initial cytoplasmic

RhoA concentration $\rho_{cp}(t=0)$ are specified as

$$R_{cp} = 1 - \sum_{i=1}^{N} R_a^i - \sum_{i=1}^{N} R_{in}^i \,, \tag{S4a}$$

$$\rho_{cp} = 1 - \sum_{i=1}^{N} \rho_a^i - \sum_{i=1}^{N} \rho_{in}^i \,, \tag{S4b}$$

where $N$ is the total number of vertices in our model. The polarized Rac1 promotes mechanical polarization. Once the GTPases in the cell settle to the steady state (Fig.S3), both Rac1 and RhoA concentrations are re-initialized (see Fig.S1). These initial conditions and re-initialization procedure enable continuous and strongly directional cell migration.

In the durotaxis study, the simulation setup must ensure that the durotaxis emerges from purely mechanical cell-ECM interactions. In other words, we need to exclude the possibility that directional migration is the result of the persistently polarized Rho GTPase signals. To switch off the chemically induced polarization, we set the active Rac1 to be uniform random distribution with small concentration differences across the vertices whereas we set other types of signals as uniform distributions without randomness (Fig.S4). Then, we run $n$ number of simulations to extract the mean durotactic index $\overline{DI}$ and the mean migration speed $\overline{V}$ as well as their standard deviations.

### S4 Analytical derivation of the substrate deformation force

We calculated the substrate force $F_{sub}$ analytically and numerically at a single FA site represented by the motor-clutch model (Fig.S5 A). The isolated motor-clutch dynamics is numerically computed from Eqs. 2–6 for $N_c = N_c^0$ , $N_m = N_m^0$ , and the protrusion resistance force $F_p = 0$ , which is otherwise determined by Eq. 8. We explain our ansatz $F_p = 0$ as follows: After clutch formation, $F_p$ builds up at a much slower rate ($\sim 1/T$) than $F_{sub}$ . Therefore, the local mechano-sensitive response of the cell at an FA site, specifically the dependence of actomyosin pulling force strength $F_{am}$ on the substrate stiffness $K_{sub}$ and subsrate viscosity $\gamma_{sub}$ must predominantly be set by $F_{sub} = F_c$ (Eqs. 2, 6). To that end, the analysis of $F_{sub}$ in Fig. 3 along with the clutch complexation dynamics yields a threshold stiffness and viscosity that explain the non-monotonic behavior of the migration speed in the whole cell simulations.

The assumption $F_p = 0$ allows us to derive an analytical expression for the substrate force $F_{sub}$ by simulatenaously solving Eqs. 2, 3, 4, and 6. The substrate force starts building up at an FA site when the molecular clutches bind between the actin filaments and the substrate with a constant association rate $k_{on}$ , or equivalently at a time scale $\tau_{on} \equiv 1/k_{on}$ . At any instant, the number of the associated clutches is $n_c \equiv PN_c^0$ . As per Eq. 6, the total restoring force of the bounded clutches is balanced by the substrate deformation. For an elastic substrate, we have a simple force-displacement relation,

$$K_{sub} x_{sub} = n_c K_c \left( x_r - x_{sub} \right) \,. \tag{S5}$$

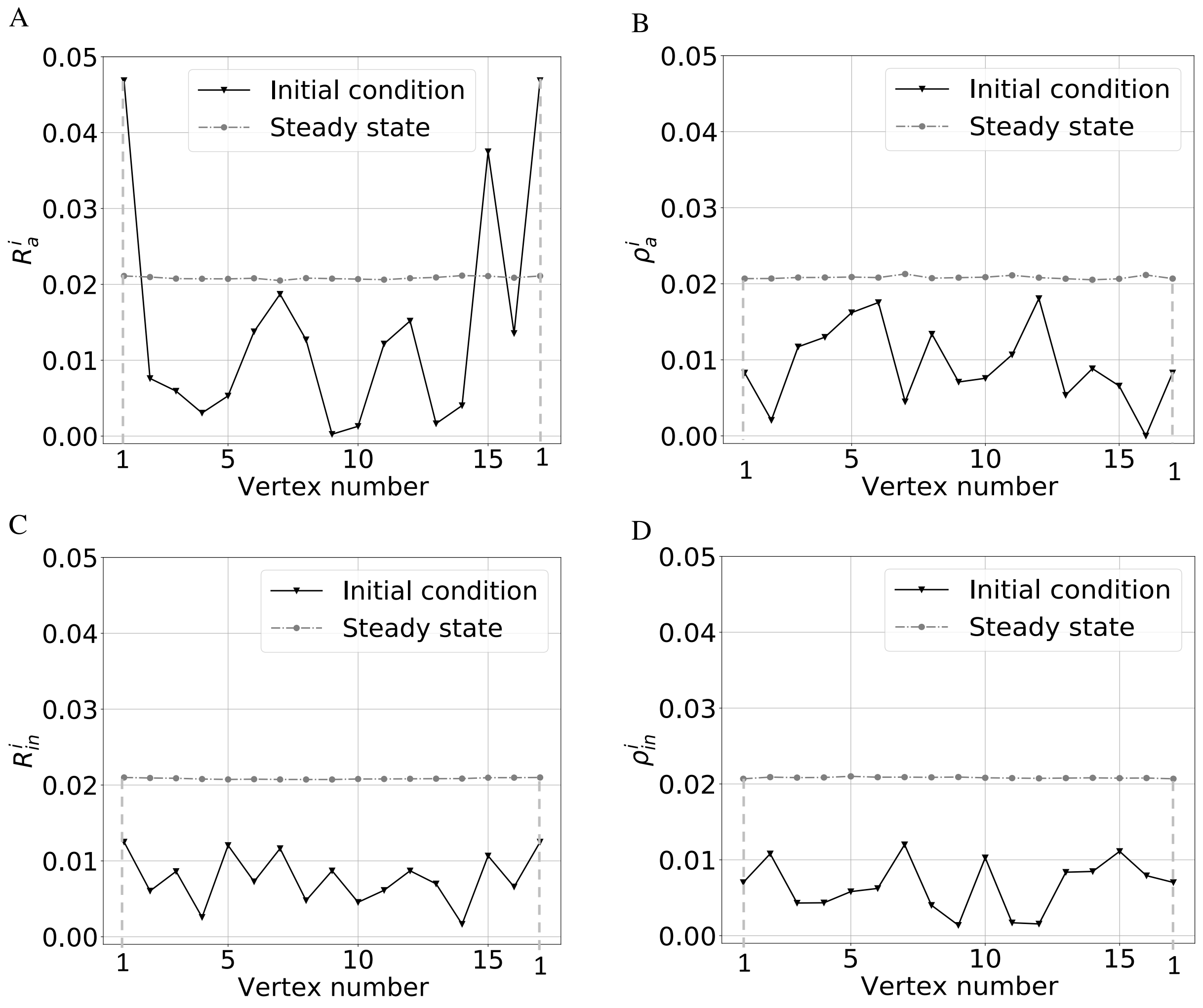


Fig. S3: **Initial conditions of the membrane-bound Rac1 and RhoA signals for migration on uniform substrates.** (A) The representative initial condition of the active Rac1 signal $R_a^i$ across all vertices ($N = 16$). The corresponding steady state is also shown. (B) Randomly distributed initial active RhoA signal $\rho_a^i$ and its steady state. (C-D) Random distributions of the inactive Rac1 signal $R_{in}^i$ and the inactive RhoA signal $\rho_{in}^i$ and their steady states.

The time derivative of Eq.S5 together with the force-velocity relation $\dot{x}_r = V_0 \left(1 - \frac{K_{sub} x_{sub}}{N_m^0 f_m}\right)$ gives

$$\dot{x}_{sub} = \frac{n_c K_c V_0 K_{sub}}{N_m^0 f_m (K_{sub} + n_c K_c)} \left( \frac{N_m^0 f_m}{K_{sub}} - x_{sub} \right) , \tag{S6}$$

assuming that the bound clutch fraction is saturated at $P \sim 1$ . The substrate displacement starting from

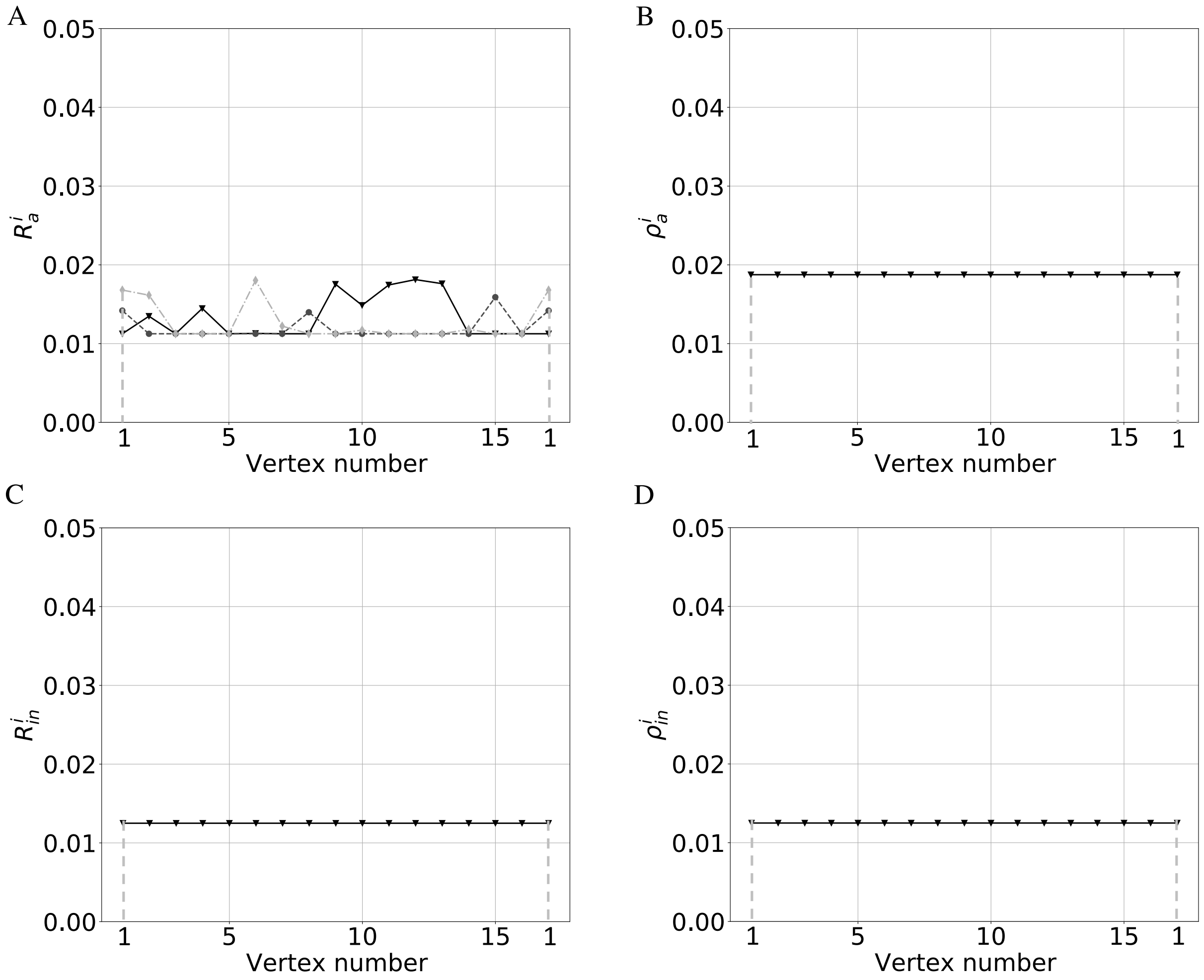


Fig. S4: **Initial conditions of the membrane-bound Rac1-RhoA signals for migration on substrates with stiffness gradients.** (A) Randomly distributed Rac1 signal. (B-D) All other types of membrane-bound proteins, i.e., $\rho_a^i$, $R_{in}^i$, $\rho_{in}^i$, are uniformly distributed on vertices.

a value of zero at $t = 0$ can be given by the solution of Eq. S6,

$$x_{sub} = \frac{N_m^0 f_m}{K_{sub}} \left(1 - e^{-t/\tau_l}\right) , \tag{S7}$$

where $\tau_l \equiv \frac{N_m^0 f_m}{V_0 K_{sub}} + \frac{N_m^0 f_m}{n_c K_c V_0}$. The time scale $\tau_l$ sets the substrate deformation rate. For compliant substrates ($K_{sub} \ll N_c^0 K_c$), the substrate deformation time scale can be approximated as $\tau_l \approx \frac{N_m^0 f_m}{V_0 K_{sub}}$. At $t = \tau_l$, substrate force is sufficiently developed, and the fraction of the bounded clutches is maximum. At $t > \tau_l$, the clutch disassociation rate $k_{off}$ rapidly increases, leading to the quick rupture of the bounded clutches. It follows that the constant $\tau_l$ can qualitatively estimate the lifetime of a binding–unbinding cycle $\tau_{off}$ in the FA dynamics (Fig.S5 B). Fig.S5C indicates a good agreement between $\tau_l$

and $\tau_{off}$ from the simulations. Based on these results, on soft substrates ($\tau_l > \tau_{on}$), a large number of clutches form, i.e., $n_c \sim N_c^0$ . In contrast, stiff substrates will induce a short lifetime of a FA cycle ($\tau_l < \tau_{on}$). As a result, only limited fractions of clutches can bind to the substrate during that short time, contributing to a low substrate traction force. In light of this, by comparing the binding time scale $\tau_{on}$ to the substrate deformation time scale $\tau_l$, we obtain a threshold stiffness,

$$K_0 = \frac{n_m f_m k_{on}}{V_0} \,. \tag{S8}$$

With this threshold stiffness, we classify the substrate as soft if $K_{sub}/K_0 < 1$ and stiff if $K_{sub}/K_0 > 1$ .

For a viscoelastic substrate, the substrate force-displacement relation is given by

$$\gamma_{sub}\dot{x}_{sub} + K_{sub}x_{sub} = n_c K_c \left(x_r - x_{sub}\right) \,. \tag{S9}$$

Differentiating Eq.S9 with respect to the time $t$ and substituting the force-velocity relation $\dot{x}_r = V_0 \left(1 - \frac{\gamma_{sub}\dot{x}_{sub} + K_{sub}x_{sub}}{N_m^0 f_m}\right)$ gives,

$$\frac{\gamma_{sub}}{n_c K_c}\ddot{x}_{sub} + \left(1 + \frac{K_{sub}}{n_c K_c} + \frac{V_0 \gamma_{sub}}{N_m^0 f_m}\right)\dot{x}_{sub} + \frac{V_0 K_{sub}}{N_m^0 f_m} x_{sub} = V_0 \,. \tag{S10}$$

On compliant substrates ($K_{sub} \ll n_c K_c$) the time scale separation $\gamma_{sub}/n_c K_c \ll \tau_l$ occurs for the range of substrate viscosity $\gamma_{sub}$ we consider in the simulations (Table 1, S2). Then, Eq.S10 is simplified to

$$\left(1 + \frac{V_0 \gamma_{sub}}{N_m^0 f_m}\right)\dot{x}_{sub} + \frac{V_0 K_{sub}}{N_m^0 f_m} x_{sub} = V_0 \,. \tag{S11}$$

Solving the equation with the initial condition $x_{sub}(t = 0) = 0$, we obtain,

$$x_{sub} = \frac{N_m^0 f_m}{K_{sub}} \left[1 - \exp\left(-\frac{t}{(\tau_l + \tau_r)}\right)\right] \,, \tag{S12}$$

where $\tau_r \equiv K_{sub}/\gamma_{sub}$ is the retardation time scale of the Kelvin-Voigt model. The total substrate force is then given by,

$$F_{sub} = \gamma_{sub}\dot{x}_{sub} + K_{sub}x_{sub} = N_m^0 f_m \left[1 - \left(\frac{\tau_l}{\tau_l + \tau_r}\right) \exp\left(-\frac{t}{(\tau_l + \tau_r)}\right)\right] \,. \tag{S13}$$

The time scale $\tau_l$ still provides a qualitative estimation of the lifetime of the clutch binding–unbinding cycle for soft substrates with sufficiently low viscosities (Fig.S5 C). On a very viscous substrate, the retardation time scale $\tau_r$ increases suppressing substrate displacement, which rapidly increases the restoring force of the bounded clutches. As a result, the force-dependent clutch disassociation rate becomes much larger than the association rate ($k_{off} \gg k_{on}$), and the bounded clutches rupture with a short lifetime (see Fig. 3F). As shown in Fig. S5 D, there is a threshold viscosity $\gamma_0$, beyond which the clutches have a very short lifetime, in turn leading to a low substrate force. We determine the threshold viscosity $\gamma_{sub}$ by locating the sudden drop of the lifetime as a function of the substrate stiffness $K_{sub}$ , as plotted in Fig. 3C (black line). The influence of viscosity is insignificant on stiff substrates ($K_{sub} > K_0$) since all clutches

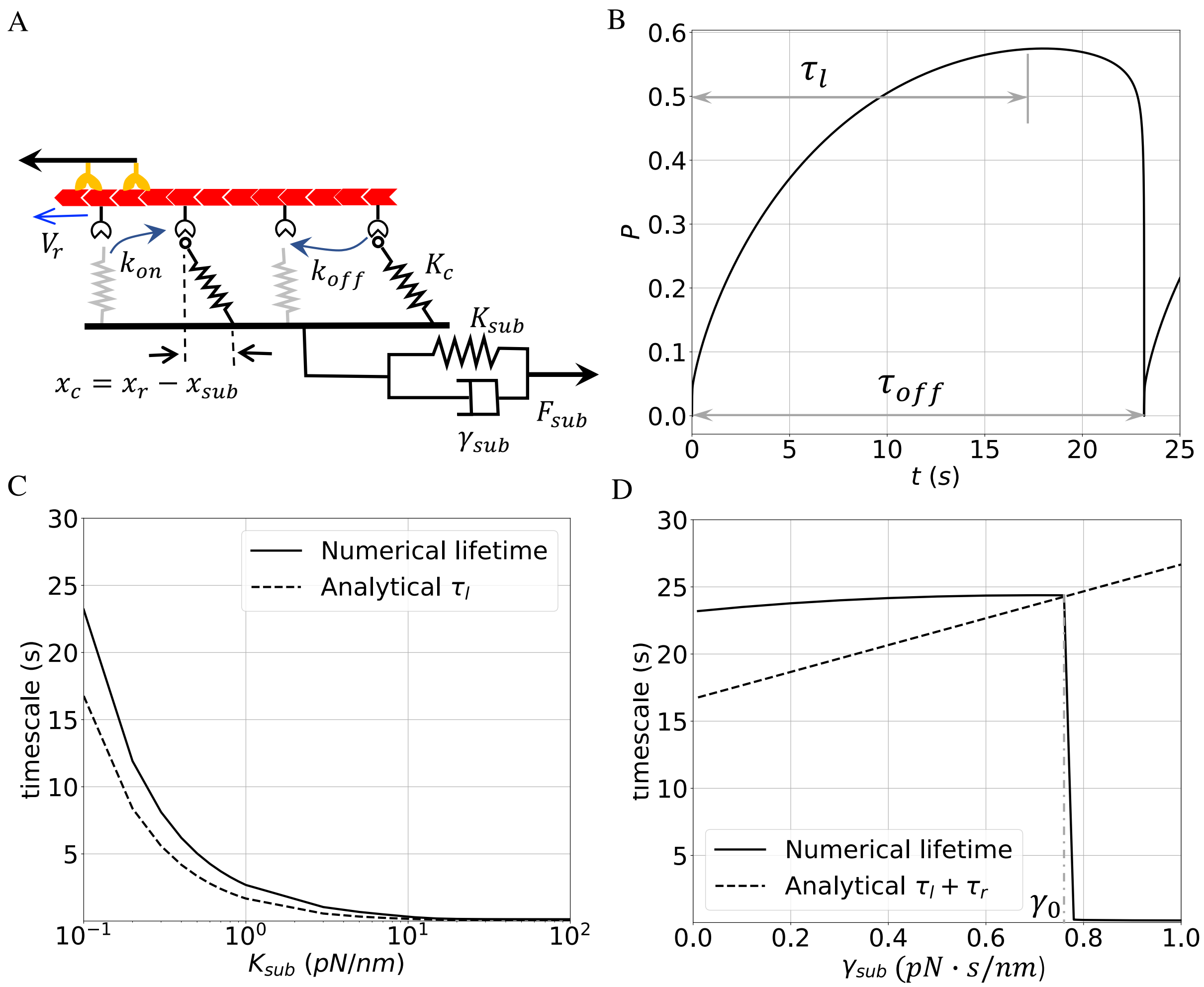


Fig. S5: **FA analysis investigating mechano-sensing at the sub-cellular level.** (A) An isolated motor-clutch model without the influence of chemical signaling and mechanical forces from the cell body. (B) Characteristic time scale $\tau_l$ provides an estimation of the life time $\tau_{off}$ of the clutch binding/unbinding cycle. (C) The lifetime of a clutch binding/unbinding cycle versus stiffness of the elastic substrate. Numerical results are obtained from single motor-clutch simulations. (D) The lifetime of an FA cycle as a function of the substrate viscosity. Substrate stiffness stays constant with $K_{sub} = 0.1\ pN/nm$. The threshold viscosity is denoted by $\gamma_0$ and corresponds to the sudden drop of the clutch lifetime.

rupture due to the condition $\tau_{on} > \tau_{off}$ prior to the viscoelastic relaxation of the substrate deformations.

### S5 Durotaxis on substrates with different gradients

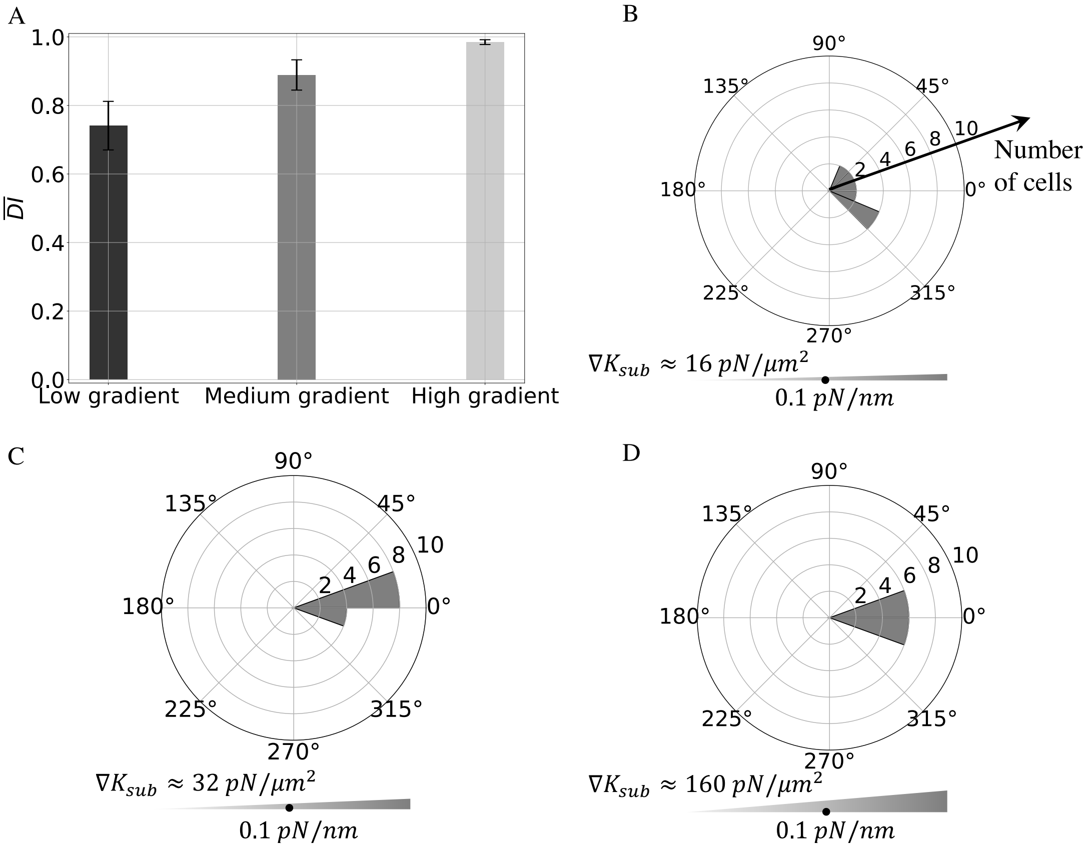


Fig. S6: **Increasing stiffness gradient enhances cell durotactic efficiency.** (A) Distributions of $\overline{DI}$ values for cells on soft elastic substrates with $K_{sub,0} = 0.1\ pN/nm$ with different gradients ($\gamma_{sub} = 0$). Low, medium, and high gradients corresponding to 16 $pN/\mu m^2$, 32 $pN/\mu m^2$, and 160 $pN/\mu m^2$, respectively. Error bars: standard deviations over $n = 12$ simulations. (B-D) Angular displacement plots for cells on different gradient substrates. The area of each sector is proportional to the number of cells moving in the corresponding direction.
