## Supplementary figures and images for "A multiscale whole-cell theory for mechano-sensitive migration on viscoelastic substrates"

### Movie1

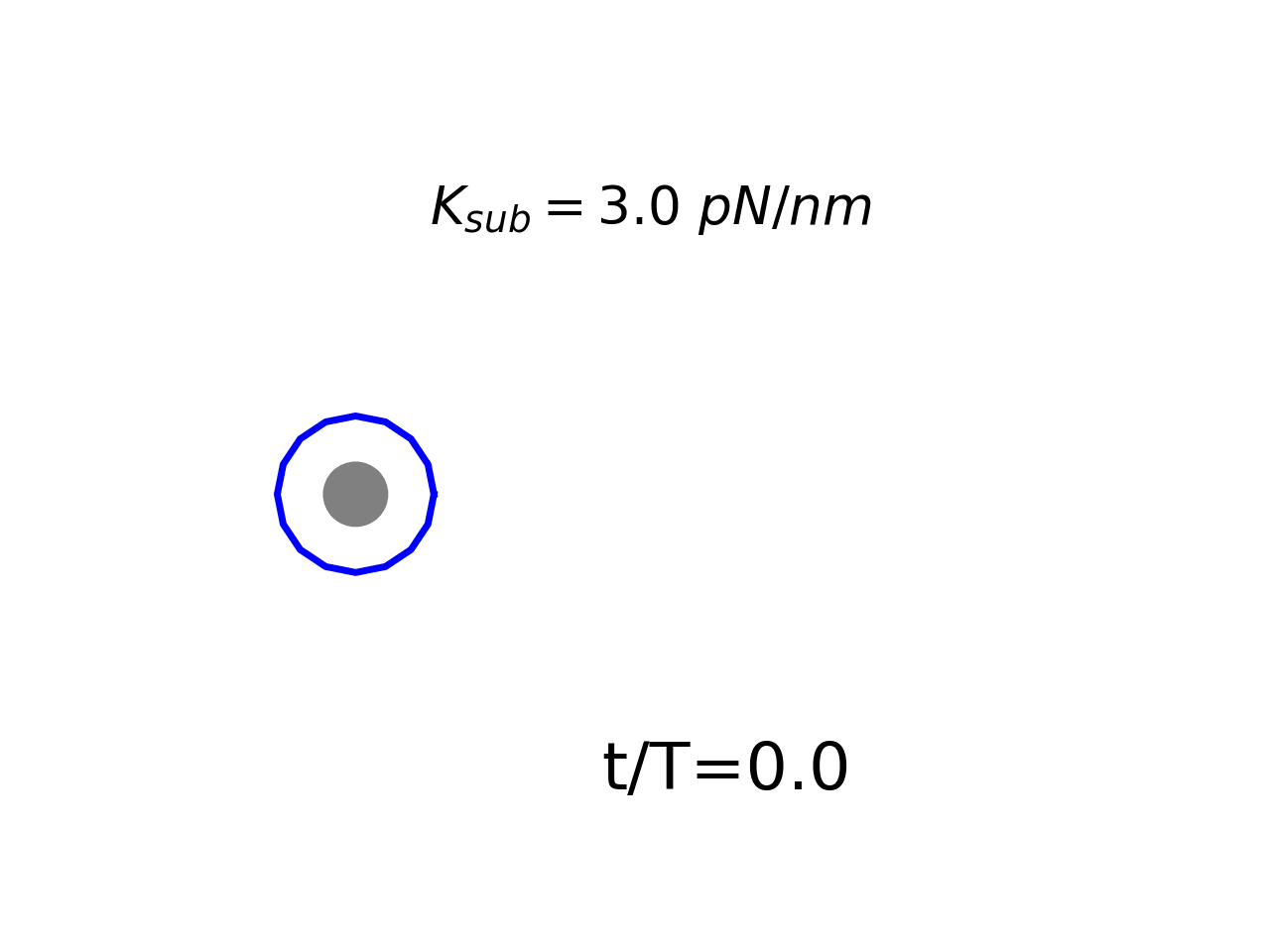

### Movie2

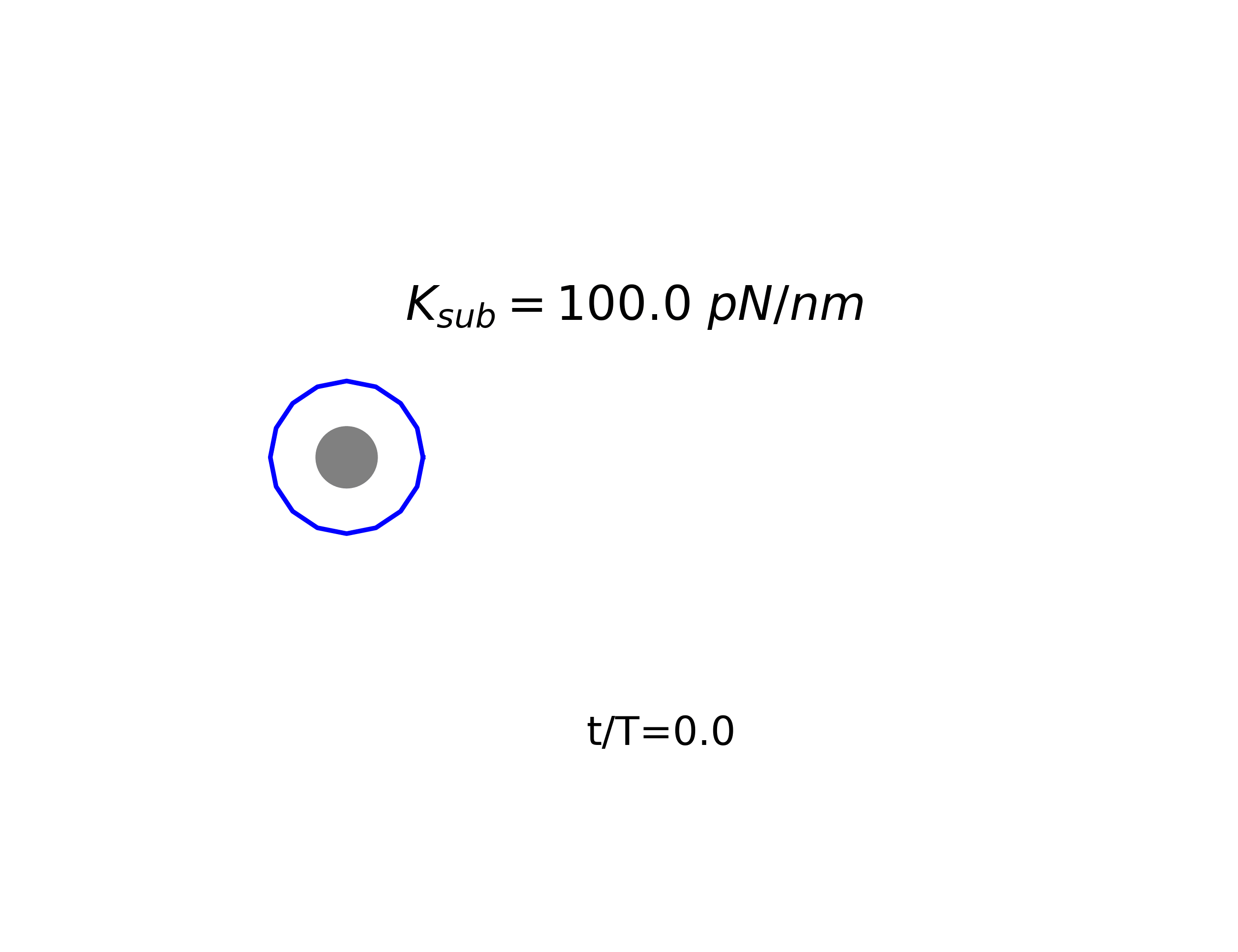
